# OpenNucleome for high resolution nuclear structural and dynamical modeling

**DOI:** 10.1101/2023.10.16.562451

**Authors:** Zhuohan Lao, Kartik Kamat, Zhongling Jiang, Bin Zhang

## Abstract

The intricate structural organization of the human nucleus is fundamental to cellular function and gene regulation. Recent advancements in experimental techniques, including high-throughput sequencing and microscopy, have provided valuable insights into nuclear organization. Computational modeling has played significant roles in interpreting experimental observations by reconstructing high-resolution structural ensembles and uncovering organization principles. However, the absence of standardized modeling tools poses challenges for furthering nuclear investigations. We present OpenNucleome—an open-source software designed for conducting GPU-accelerated molecular dynamics simulations of the human nucleus. OpenNucleome offers particle-based representations of chromosomes at a resolution of 100 KB, encompassing nuclear lamina, nucleoli, and speckles. This software furnishes highly accurate structural models of nuclear architecture, affording the means for dynamic simulations of condensate formation, fusion, and exploration of non-equilibrium effects. We applied OpenNucleome to uncover the mechanisms driving the emergence of “fixed points” within the nucleus—signifying genomic loci robustly anchored in proximity to specific nuclear bodies for functional purposes. This anchoring remains resilient even amidst significant fluctuations in chromosome radial positions and nuclear shapes within individual cells. Our findings lend support to a nuclear zoning model that elucidates genome functionality. We anticipate OpenNucleome to serve as a valuable tool for nuclear investigations, streamlining mechanistic explorations and enhancing the interpretation of experimental observations.

## Introduction

The highly complex structural organization of the human nucleus plays a crucial role in the functioning and regulation of our cells.^1–10^ The complexity arises from the diverse range of nuclear landmarks, such as nucleoli,^11^ nuclear speckles,^8,12^ and the nuclear lamina,^13^ each serving distinct functions. These landmarks provide specialized environments for various nuclear processes, allowing for efficient coordination and regulation of gene expression. Moreover, the spatial arrangement of chromosomes within the nucleus, intertwined with the nuclear landmarks, is critical for proper gene regulation and communication between different genome regions. Disruptions or abnormalities in the nuclear organization can have profound consequences on cellular function and can contribute to the development of diseases, including cancer and genetic disorders.^14,15^

Recent advancements in experimental techniques have significantly enhanced our understanding of nuclear organization.^3,16–20^ The advent of high-throughput sequencing-based methods, such as genome-wide chromosome-conformation capture (Hi-C), has unveiled crucial structural elements of the genome,^21,22^ including chromatin loops,^23^ topologically associating domains,^24,25^ and compartments.^22^ Additionally, sequencing-based techniques such as DamID,^26^ Chip-Seq,^27^ and TSA-Seq^28^ have revealed valuable information regarding interactions between chromosomes and nuclear landmarks. However, it is worth noting that these sequencing methods often offer averaged contacts, which can mask the heterogeneity present across populations, although single-cell techniques are also emerging.^29–31^ Moreover, translating contact data into spatial positions can be challenging, adding complexity to interpreting experimental findings.

To complement these sequencing approaches, microscopic imaging techniques directly probe the spatial positions within individual nuclei.^3,13,32–34^ Recent advancements in DNA FISH (Fluorescence In Situ Hybridization) have enabled high-throughput imaging of thousands of loci simultaneously.^35,36^ These imaging studies have not only confirmed the structural features observed through sequencing techniques but have also provided valuable insights into the heterogeneity present at the single-cell level.

The abundance of available experimental data in the field of nuclear organization provides a fertile ground for structural modeling.^37–63^ To make sense of this wealth of information, various computational approaches have been introduced, with polymer simulation approaches being extensively utilized. These simulation techniques aid in reconstructing structural ensembles that closely replicate experimental data, offering valuable insights into the mechanisms underlying chromosome folding. In recent studies, these approaches have also been employed to investigate the interplay between the genome and the nuclear lamina, ^64–67^ as well as nucleoli,^68^ shedding light on their dynamic relationships.

Despite the progress made in computational modeling, the absence of well-documented software with easy-to-follow tutorials pose a challenge. Many research groups develop their own independent software, which complicates cross-validation and hinders the establishment of best practices for genome modeling. ^40,69,70^ Moreover, comprehensive models of the entire nucleus, especially at high resolution, remain scarce. Addressing these limitations and fostering collaboration in the scientific community can be achieved through the development of open-source tools. By promoting transparency and accessibility, such tools have the potential to greatly facilitate nuclear modeling and contribute to a more unified and collaborative research environment.

We present OpenNucleome, an open-source software designed for conducting molecular dynamics (MD) simulations of the human nucleus. This software streamlines the process of setting up whole nucleus simulations through just a few lines of Python scripting. OpenNucleome can unveil intricate, high-resolution structural and dynamic chromosome arrangements at a 100 KB resolution. It empowers researchers to track the kinetics of condensate formation and fusion while also exploring the influence of chemical modifications on condensate stability. Furthermore, it facilitates the examination of nuclear envelope deformation’s impact on genome organization. The software’s modular architecture enhances its adaptability and extensibility. Leveraging the power of OpenMM,^71^ a GPU-accelerated MD engine, OpenNucleome ensures efficient simulations.

Our work demonstrates the fidelity of the simulated nuclear organizations by faithfully reproducing Hi-C, Lamin B DamID, TSA-Seq, and DNA-MERFISH data. The dynamic insights extracted from this model are pivotal in advancing our understanding of nuclear organization mechanisms. Our findings reveal that inherent heterogeneity in chromosome contacts naturally emerges within single cells. Interestingly, robust contacts between chromosomes and nuclear bodies can also be established due to a coupled self-assembly mechanism. Notably, the resilience of contacts involving nuclear bodies supports a nuclear zoning model for genome function. In the realm of nuclear investigations, we anticipate OpenNucleome to serve as an invaluable tool, seamlessly complementing experimental techniques.

## Results

### Non-equilibrium nucleus model at 100 KB resolution

We present an open-source implementation of a computational framework that facilitates the structural and dynamical characterization of the human nucleus. This framework builds upon a previous investigation^65^ but incorporates several significant modifications. Firstly, we enhance the model resolution by a factor of ten, enabling the precise determination of the spatial positioning of each chromatin segment measuring 100 KB in length. Secondly, we present a kinetic scheme for speckles that accounts for the phosphorylation of protein molecules. This inclusion captures the influence of chemical reactions on the stability and dynamics of nuclear bodies. Thirdly, we incorporate explicit nuclear envelope dynamics to explore the impact of large-scale deformations on genome organization. Finally, our implementation into OpenMM offers the advantages of Python Scripting and GPU acceleration, facilitating easy extension and customization. These features will facilitate the broad applicability and adoption of the proposed model.

The nucleus model provides particle-based representations for chromosomes, nucleoli, speckles, and the nuclear envelope. As shown in Fig. 1A and B, each of the 46 chromosomes is represented as a beads-on-a-string polymer, where each bead represents a 100-KB-long genomic segment. Based on Hi-C data, we further assign each bead as compartment *A, B,* or *C* to signify euchromatin, heterochromatin, or pericentromeric regions. The lamina was modeled as a spherical enclosure with 10 *µm* diameter, using discrete particles arranged to represent a mesh grid with covalent bonds linking together nearest neighbors.^72^ We modeled nucleoli and speckles as liquid droplets that emerge through the spontaneous phase separation of coarse-grained particles, representing protein and RNA molecule aggregates.^8,11^ These particles exhibited attractive interactions within the same type to promote condensation. More details about the various components of the system can be found in the Supporting Information *Section: Components of the whole nucleus model*.

**Figure 1:**
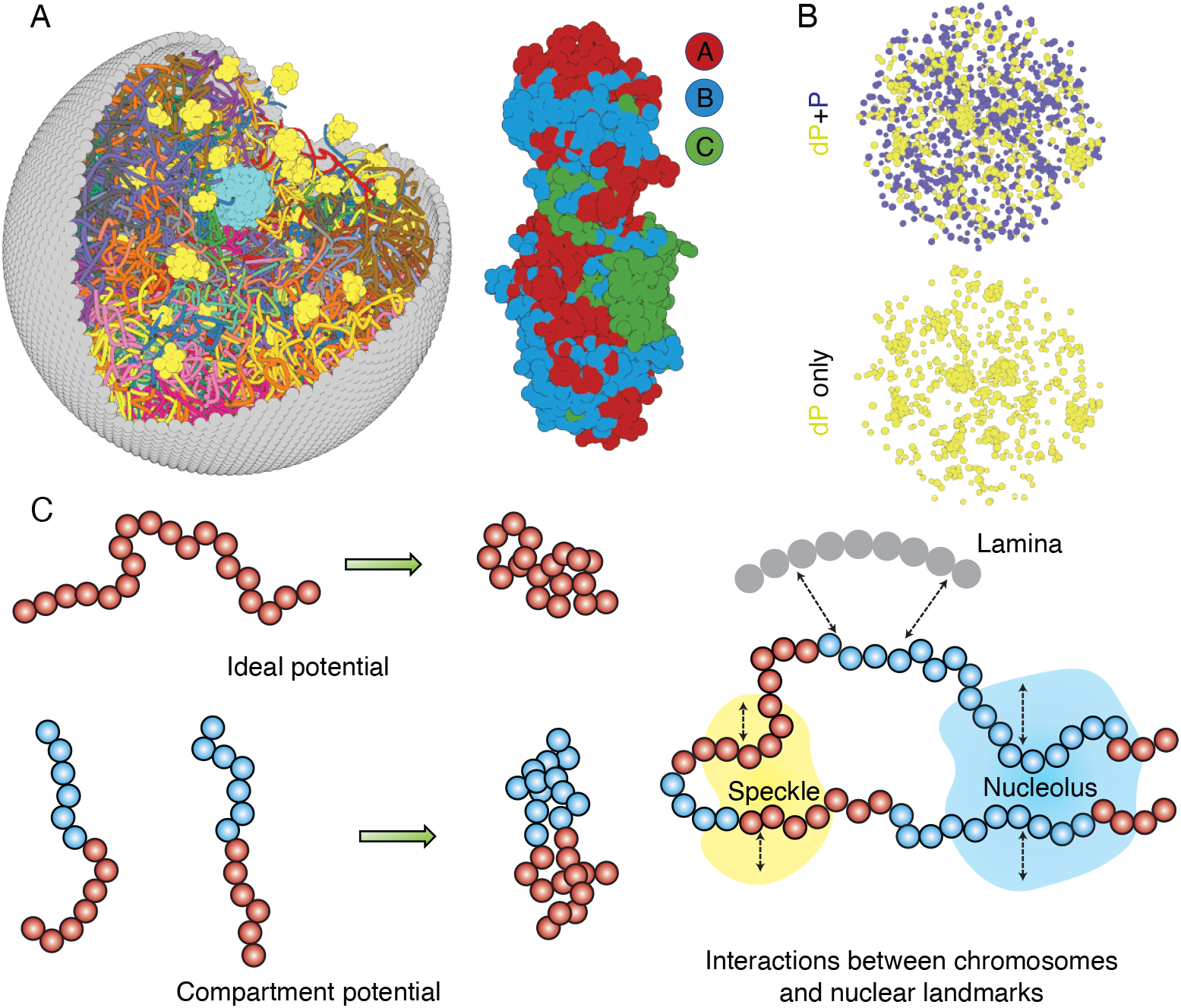
Computer model of the human nucleus for structural and dynamical characterizations. (A) 3D rendering of the nucleus model with particle-based representations for the 46 chromosomes shown as ribbons, the nuclear lamina (grey), nucleoli (cyan), and speckles (yellow). As shown on the right, chromosomes are modeled as beads-on-a-string polymers at a 100 KB resolution, with the beads further categorized into compartment A (red), compartment B (light blue), or centromeric regions (green). (B) Speckle particles undergo chemical modifications concurrent to their spatial dynamics, and the dephosphorylated (dP) particles contribute to droplet formation. (C) Illustration of the ideal and compartment potential that promotes chromosome compaction and microphase separation. Specific interactions between chromosomes and nuclear landmarks are shown on the right.

The energy function of the nucleus model includes three components that account for the self-assembly of chromosomes, the assembly of nuclear bodies, and the coupling between chromosomes and nuclear landmarks. Therefore, the model approximates nuclear organization as a coupled self-assembly process. The chromosome energy function (see Eq. S7 in Supporting Information *Section: Hi-C inspired interactions for the diploid human genome*) includes terms that account for the polymer connectivity and excluded volume effect, an ideal potential, compartment-specific interactions, and specific interchromosomal interactions. As shown in Fig. 1C, the ideal potential is only applied for beads from the same chromosome to approximate the effect of loop extrusion by Cohesin molecules ^73,74^ for chromosome compaction and territory formation.^75,76^ Compartment-specific interactions, on the other hand, promote microphase separation and compartmentalization of euchromatin and heterochromatin. Finally, interchromosomal interactions account for sequence-specific effects that compartment-dependent potentials cannot capture.

Interactions among coarse-grained particles that form nuclear bodies were designed to promote and stabilize the formation of liquid droplets, as has been revealed by many experiments.^77–79^ We adopted the Lennard-Jones potential for nucleolar particles to mimic the weak, multivalent interactions that arise from protein and RNA molecules that make up the nucleoli. As a first attempt to approximate their complex dynamics, we considered two types of particles that form speckles: phosphorylated (P) and de-phosphorylated (dP). The two types can interconvert via chemical reactions^80–82^ and dP particles share attractive interactions modeled with the Lennard-Jones potential.

As shown in Fig. 1C, to recognize specific interactions between chromosomes and nuclear landmarks, we introduced contact potentials between them. These potentials are inspired by the experimental techniques that probe the corresponding contacts. The Supporting Information *Section: Chromosome-nuclear landmark interactions* and *Nuclear landmark-nuclear landmark interactions* contain more details about all the nuclear landmark related energy functions.

### Optimization of model parameters with experimental data

The nucleus model was designed to be interpretable such that energy terms represent physical processes. Furthermore, the expressions of the interaction potentials were also designed such that their parameters can be determined from experimental data via the maximum entropy optimization algorithm.^9,83,84^ Below, we briefly outline the procedure used for parameter optimization and further details can be found in the Supporting Information *Section: Optimization of the whole nucleus model parameters*.

As illustrated in Fig. 2A, starting from a given set of parameters, we first perform MD simulations to produce a collection of 3D structures for the diploid genome and various nuclear bodies. These structures are then transformed into a contact map or contact probabilities between chromatin beads and nuclear landmarks by averaging over homologous chromosomes. Constraints corresponding to different energy terms could be obtained from the simulated results and compared with those estimated from Hi-C, SON TSA-Seq, and Lamin B DamID profiles. Finally, the model parameters were updated based on the difference between simulated and experimental constraints using the adaptive moment estimation (Adam) optimization algorithm.^85^ The three steps can be repeated with updated parameters to improve the simulation-experiment agreement further.

**Figure 2:**
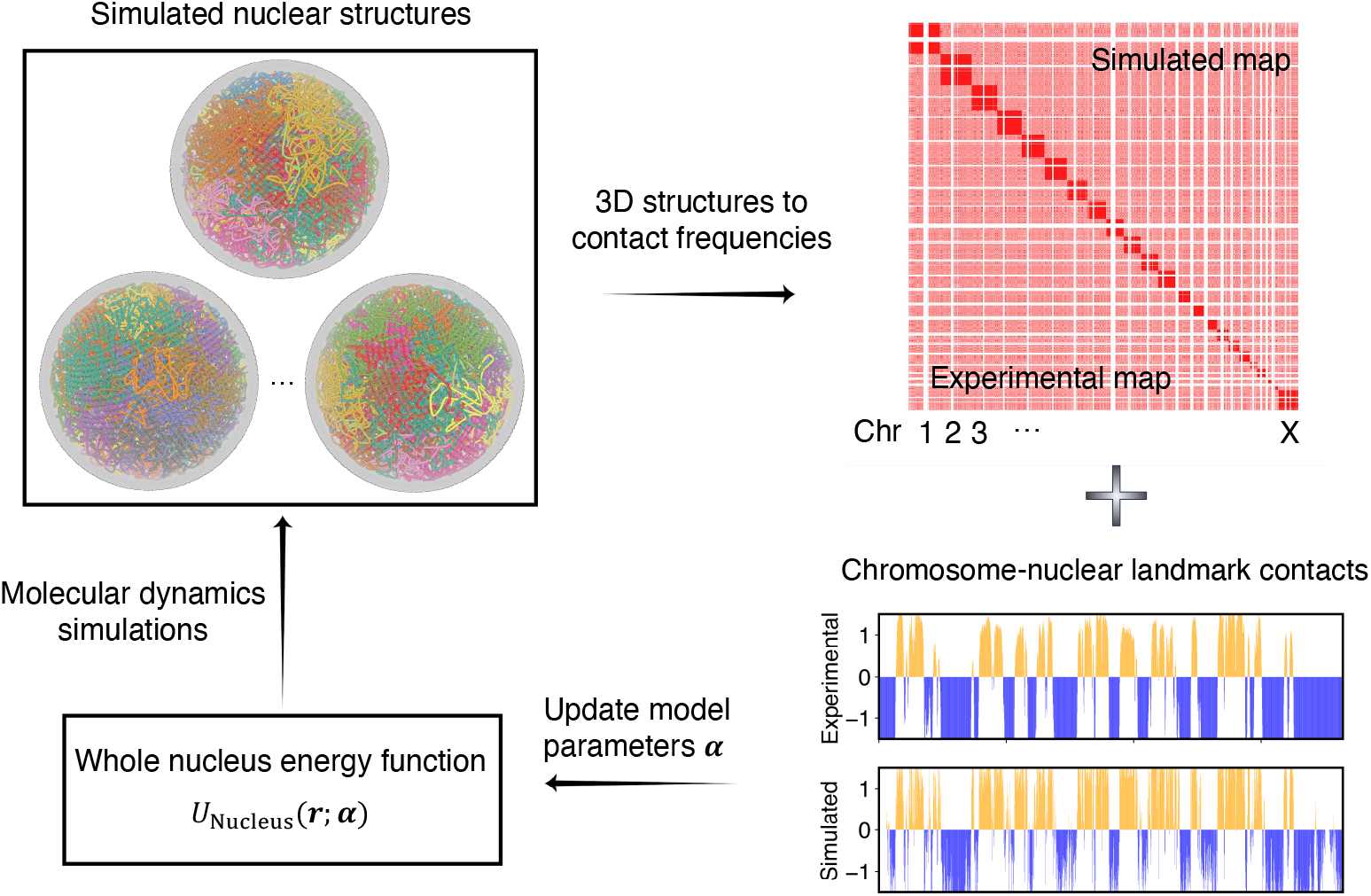
Overview of the iterative algorithm for parameterizing the nucleus model with experimental data. Starting from an initial set of parameters, we perform MD simulations to produce an ensemble of nuclear structures. These structures can be transformed into contacts between chromosomes or between chromosomes and nuclear landmarks for direct comparison with experimental data. Differences between simulated and experiment contacts are used to update parameters for additional rounds of optimization if needed.

No quantitative experimental data exists for interactions among nuclear body particles to serve as constraints. We varied the strength of the interaction potential to produce 2-3 nucleoli and ∼ 30 speckle clusters during the simulations (Fig. S1) while ensuring the fluidity of the resulting droplets.

### Molecular dynamics simulations with GPU acceleration

We implemented the nucleus model into the MD engine OpenMM.^71^ OpenMM offers an excellent interface with Python scripting, significantly improving the readability and customizability of the model. The code was designed into functional modules, with different components, such as chromosomes and nuclear landmarks, written as separate classes. This design further facilitates the introduction of additional nuclear components, if desired, with minimal changes to existing code. We provide examples of simulation set up, trajectory analysis, parameter optimization, and introducing new features in the GitHub repository.

Fig. 3A illustrates the workflow for setting up and executing whole nucleus simulations. A configuration file that provides the position of individual particles in the PDB file format is needed to initialize the simulations. This file also contains topological information regarding whether a particle represents chromosomes or nuclear landmarks and the identity of specific chromosomes. The input file can be generated with provided Python scripts by randomly distributing the positions of chromosomes, speckles, and nucleoli, though optimized configurations are also included in the GitHub repository. By default, the lamina particles will be uniformly placed on a sphere of 10 *µ*m in diameter. Upon parsing the configuration file, interactions among various components can be set up with optimized parameters. This step will produce an object that can be used for MD simulations. As shown in Fig. 3B, the workflow only requires a few lines of code. The package also includes analysis scripts to compute contact maps, monitor conformational dynamics, and track nuclear bodies.

**Figure 3:**
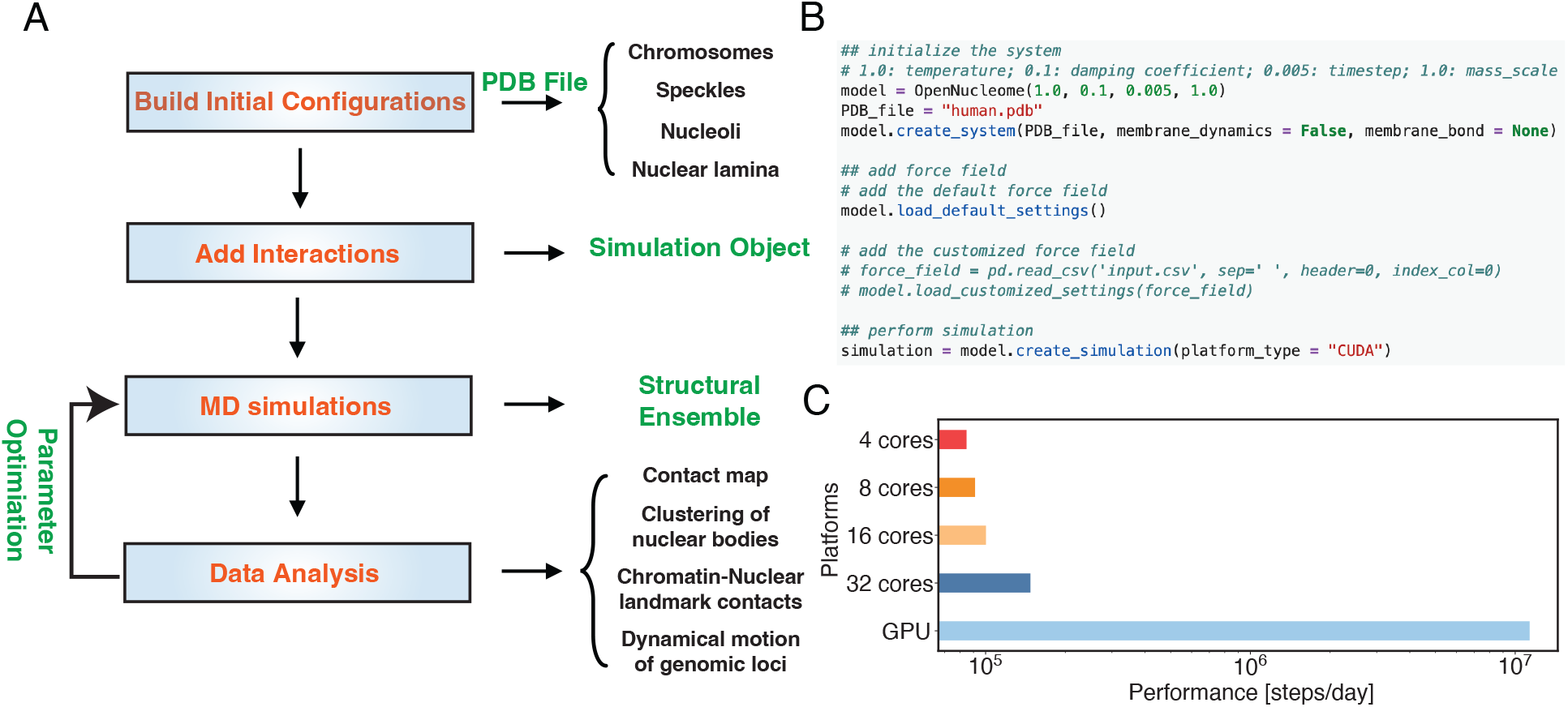
OpenNucleome facilitates GPU-accelerated simulations of the human nucleus. (A) Illustration of workflow for setting up, performing, and analyzing MD simulations. (B) Python scripts setting up whole nucleus simulations. (C) Performance of MD simulations on different number of CPU cores and a single GPU.

A significant benefit of OpenMM^71^ is its native support of GPU acceleration. As shown in Fig. 3C, the simulation speed with one Nvidia Volta V100 GPU is 150 times faster than that of the 4 Intel Xeon Platinum 8260 CPU cores. Notably, this performance enhancement cannot be achieved by simply increasing the CPU core numbers. For example, the simulation speed with 32 CPU cores is less than twice that of 4 CPU cores, potentially due to the system’s heterogeneous distribution of particles.

### Simulations reproduce and predict diverse experimental data

We extensively validated the parameterized nucleus model to examine its biological relevance. MD simulations initialized from 50 different initial configurations were performed to build an ensemble of structures. As mentioned in the following section, a diverse set of initial configurations is essential for reproducing interchromosomal contacts probed in Hi-C. From the simulated structures, we computed various quantities for direct comparison with experimental measurements. Given that the majority of experimental data were analyzed for the haploid genome, we adopted a similar approach by averaging over paternal and maternal chromosomes to facilitate direct comparison. More details on data analysis can be found in the Supporting Information *Section: Details of simulation data analysis*.

We compared the simulated contact probabilities among chromosomes with Hi-C data. As shown in Fig. 4A and Fig. S2, the simulated and experimental contact maps are highly correlated. The squares along the diagonal support the formation of chromosome territories that promote intrachromosomal contacts, and the apparent checkboard patterns follow the compartmentalization of various chromatin types. We further examined the decay of intra-chromosomal contacts as a function of the sequence separation, which is known to deviate from that of an equilibrium globule.^22^ As shown in Fig. 4B, the simulated results overlap well with the Hi-C data (orange curve). In addition, the simulated average contact probabilities between various compartment types match values estimated from Hi-C data. Moreover, the simulated and experimental average contact probabilities between pairs of chromosomes agree well, and the Pearson correlation coefficient between the two datasets reaches 0.89.

**Figure 4:**
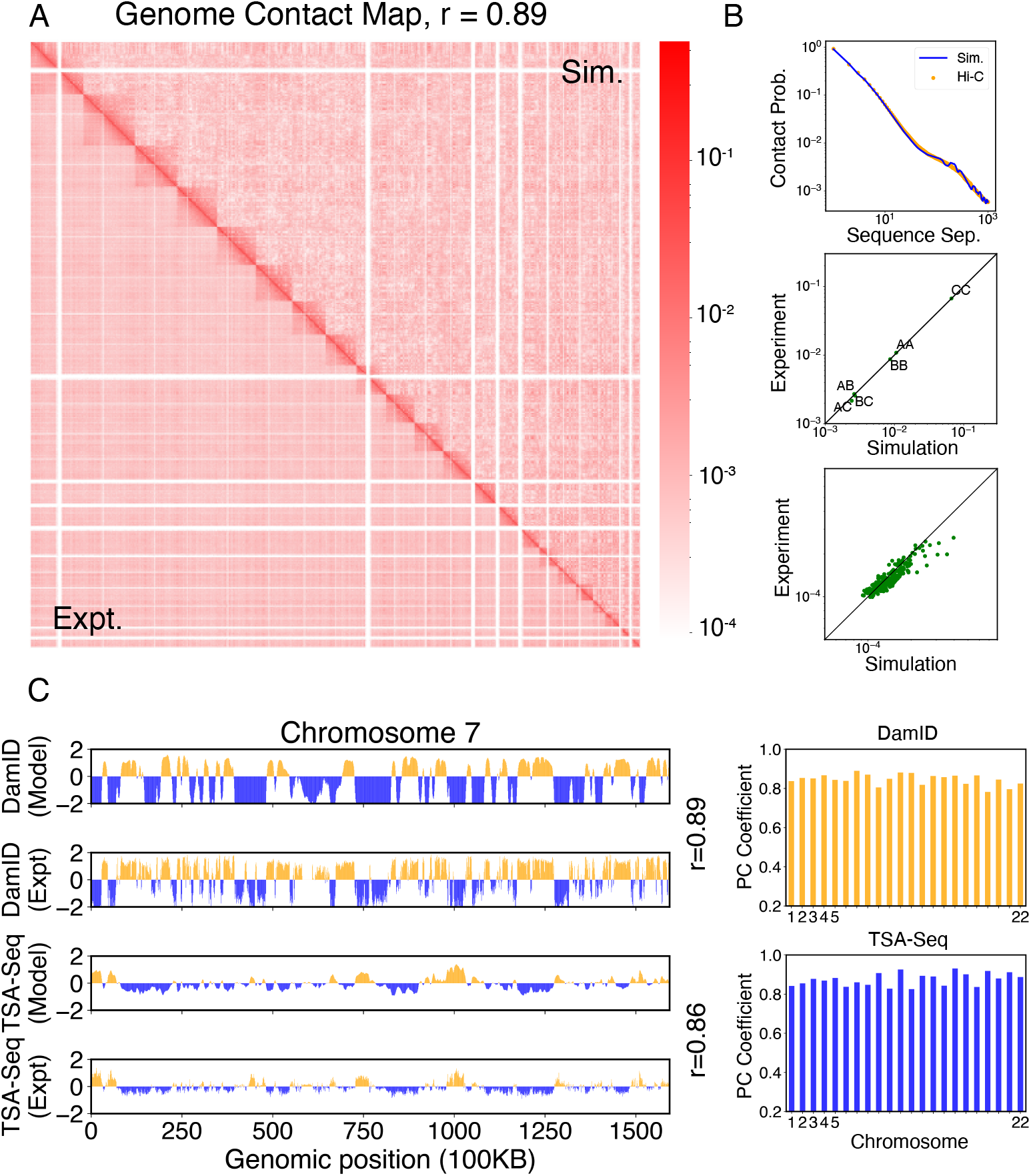
Simulated structures reproduce contact frequencies between chromosomes and between chromosomes and nuclear landmarks. (A) Comparison between simulated (top right) and experimental (bottom left) whole-genome contact probability maps with Pearson correlation coefficient *r* = 0.89. Zoom-ins of various regions are provided in Fig. S2. (B) Comparison between simulated and experimental average contact frequencies, including average contacts between genomic loci from the same chromosomes at a given separation (top), average contacts between genomic loci classified into different compartment types (middle), and average contacts between various chromosome pairs (bottom). (C) Comparison between simulated and experimental Lamin-B DamID (top) and SON TSA-Seq signals (bottom), with Pearson correlation coefficients of haploid chromosomes shown on the right.

We further examined the contacts between chromosomes and nuclear landmarks. As illustrated in Fig. 4C, the simulated Lamin-B DamID signals for chromosome 7 match well with the experimental results, capturing the complex contact pattern that weaves chromatin towards and away from the nuclear envelope. Similarly, SON TSA-Seq data that quantify the contact between chromosomes and speckles are well captured by simulated structures. The anti-correlation between DamID and TSA-Seq is clearly visible. The observed agreement between simulation and experimental results is not limited to any particular chromosome. Good agreements are achieved for all chromosomes.

The simulations also provide 3D representations of the nucleus that can be compared with DNA-MERFISH data.^35^ We found that the simulated radius of gyration of individual chromosomes matches well with experimental values (Fig. 5A). The simulated and experimental average normalized chromosome radial positions also correlate strongly, as shown in Fig. 5B. We note that while the sequencing results presented in Fig. 4 were used for model parameterization, the MERFISH data were not. Therefore, the simulation results here are *de novo* predictions, and their agreement with experimental data strongly supports the coupled assembly mechanism used for designing the energy function.

**Figure 5:**
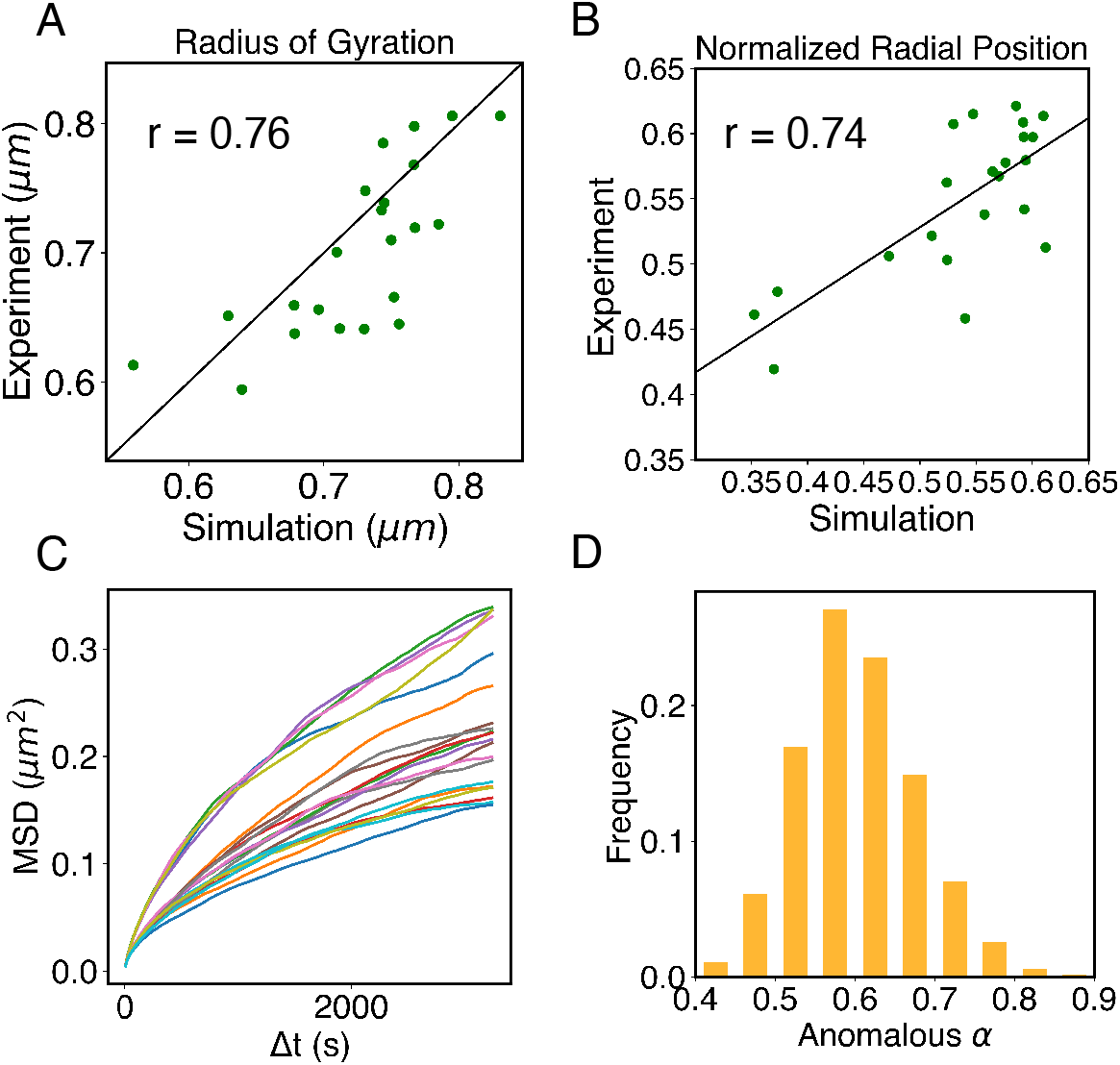
Structural and dynamical predictions of the nucleus model match results from microscopy imaging. (A) Comparison between the simulated and experimental radius of gyration, *R*_g_, for haploid chromosomes. The Pearson correlation coefficient between the two, *r*, is shown in the legend. (B) Comparison between the simulated and experimental normalized radial positions for haploid chromosomes, with their Pearson correlation coefficient shown in the legend. Detailed definition of the normalized radial positions is provided in the Supporting Information *Section: Computing simulated normalized chromosome radial positions*. (C) Mean-squared displacements, MSDs, as a function of time are shown for selected telomeres. (D) The probability distribution of the anomalous exponent, *α*, obtained from fitting the MSDs curves for all telomeres with the expression, ⟨**r**^2^(Δ*t*)⟩ = *D_α_*Δ*t^α^*.

A significant advantage of MD simulation-based models is the dynamical information they naturally produce. We measured the dynamics of telomeres by tracking the mean square displacements (MSDs), ⟨**r**^2^(Δ*t*)⟩, as a function of time. In Fig. 5C, we plot representative MSD trajectories over a one-hour timescale. In line with previous research, ^86–88^ telomeres display anomalous subdiffusive motion. When fitted with the equation ⟨**r**^2^(Δ*t*)⟩ = *D_α_*Δ*t^α^*, these trajectories yield a spectrum of *α* values, with a peak around 0.59. The exponent and the diffusion coefficient *D_α_* = (27±11)×10*^−^*^4^*µm*^2^·*s^−α^* both match well with the experimental values,^89,90^ upon setting the nucleoplasmic viscosity as 1*Pa* · *s* (see Supporting Information *Section: Mapping the reduced time unit to real time* for more details).

The good agreement in the dynamics of individual loci further inspired us to examine the diffusion of whole chromosomes. In particular, we plotted the normalized chromosome radial positions as a function of time in Fig. 6A. Remarkably, we found that chromosomes appear arrested and no significant changes in their positions are observed over timescales comparable to the cell cycle (see also Fig. S12). Therefore, our simulations predict that large-scale movements of chromosomes are unlikely during the G1 phase.

**Figure 6:**
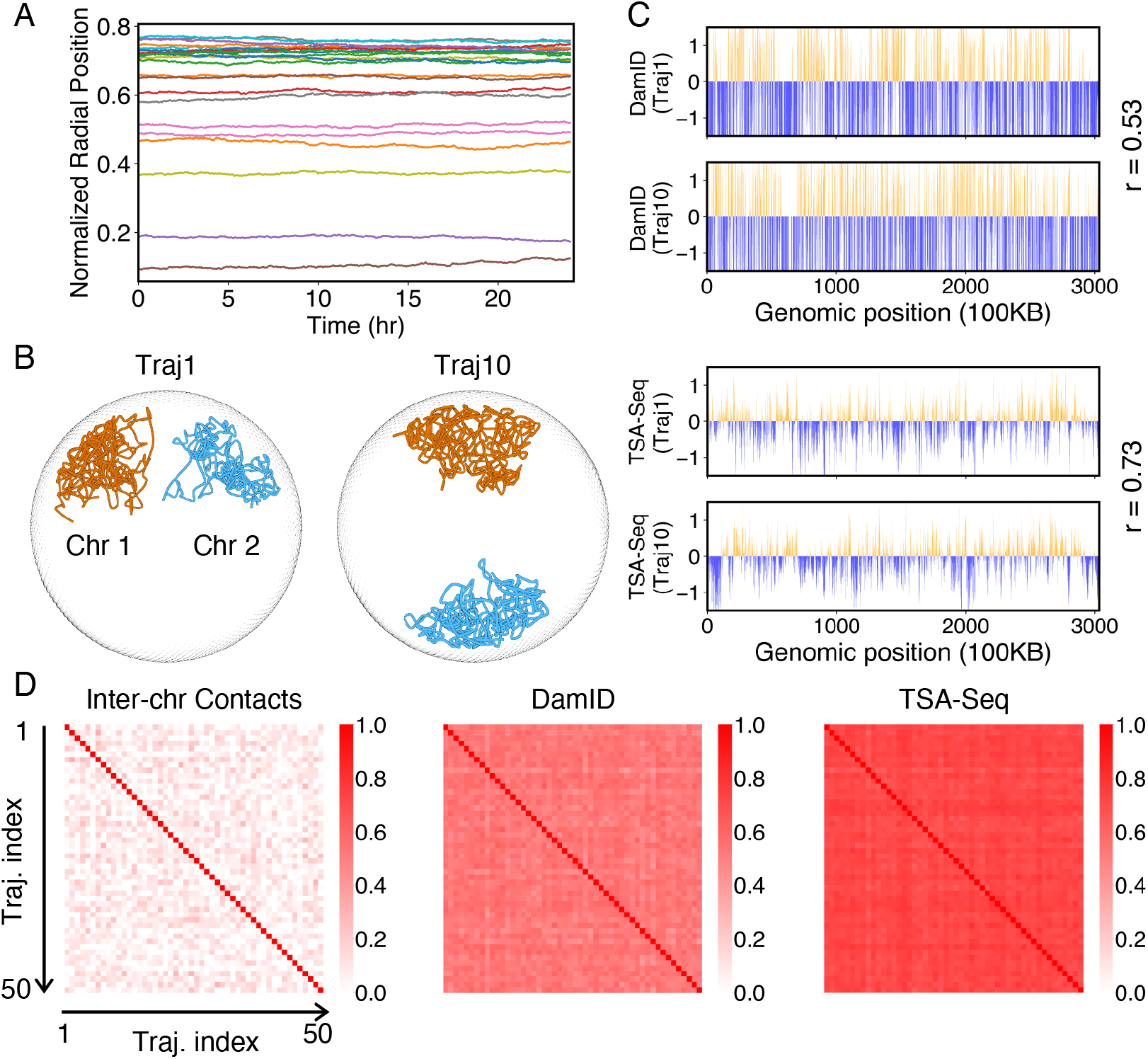
Heterogeneity and conserved features of nuclear organizations. (A) Normalized chromosome radial positions as a function of simulation time. (B) Contacts between chromosome 1 and 2 from two independent simulation trajectories show significant variations. (C) Genome-wide *in silico* Lamin B DamID (top) and SON TSA-Seq (bottom) profiles computed from two independent trajectories. Pearson correlation coefficients, r, are provided on each plot. (D) Pairwise Person correlation coefficients between interchromosomal contact matrices (left), genome-wide Lamin B DamID profiles (middle), and genome-wide SON TSA-Seq profiles (right) determined from independent trajectories. The averages excluding the diagonals of the three datasets are 0.06, 0.53, and 0.72.

### Heterogeneity and robustness of the simulated conformational ensemble

The lack of relaxation of chromosome radial positions suggests the importance of starting configurations used to initialize the simulations. Statistical averages of the resulting ensemble of nuclear structures depend crucially on these starting configurations. Using an optimization procedure, we selected them from 1000 configurations to maximize the agreement with experimental lamin-B DamID and interchromosomal contact probabilities. The Supporting Information *Section: Initial configurations for simulations* provides more details on preparing the 1000 initial configurations.

We selected a total of 50 starting configurations to initiate independent simulations. Smaller sets of starting configurations are not sufficient to reproduce the interchromosomal contact probabilities, as shown in Fig. S3B. Notably, different sets of 50 configurations selected from independent trials show significant overlap (Fig. S3D), supporting the robustness of the selection protocol in detecting conserved features of genome organization.

While the ensemble as a whole is relatively robust, individual configurations with the ensemble exhibit significant differences. For example, the Lamin B DamID profiles produced from different trajectories are only weakly correlated (Fig. 6C), with an average correlation coefficient of 0.53. These weak correlations result from significant differences in the normalized radial positions of chromosomes, as can be seen in representative configurations from two simulation trajectories (Fig. 6B). The fluctuations of normalized radial positions cause changes in contacts between chromosomes as well, resulting in little correlation between interchromosomal contact matrices (Fig. 6D).

We examined genome organizations reported by Su et al. ^35^ and found a similar variation of interchromosomal contact probabilities across individual cells (Fig. S3A, D). Notably, the simulated configurations capture the fluctuations of interchromosomal contacts observed in DNA-MERFISH data, further supporting the biological relevance of the reported *in silico* structures.

Despite the differences in interchromosomal contacts across trajectories, high conservation of connections between chromosomes and speckles can be observed in individual simulations. For example, the average correlation coefficient between *in silico* SON TSA-Seq profiles produced from different trajectories is 0.72, much higher than the corresponding value for Lamin B DamID profiles. Conservation of contacts between chromosomes and nuclear bodies (zones) across individual cells has indeed been reported in a previous study that simultaneously images chromatin and various subnuclear structures.^36^

### Nuclear deformation preserves chromosome-nuclear body contacts

Numerous studies have highlighted the remarkable influence of nuclear shape on the positioning of chromosomes and the regulation of gene expression.^53,91^ The nucleus, once regarded as a mere compartment for DNA storage, is increasingly recognized as a dynamic and intricately structured organelle. To better understand the interplay between nuclear shape and genome organization as a fundamental mechanism that shapes the transcriptional landscape, we performed additional simulations in which the nuclear lamina was altered from a sphere into more ellipsoidal shapes by applying a force along the *z*-axis (Fig. 7A). More details about these simulations can be found in the *Section: Nuclear envelope deformation simulations* of the Supporting Information.

**Figure 7:**
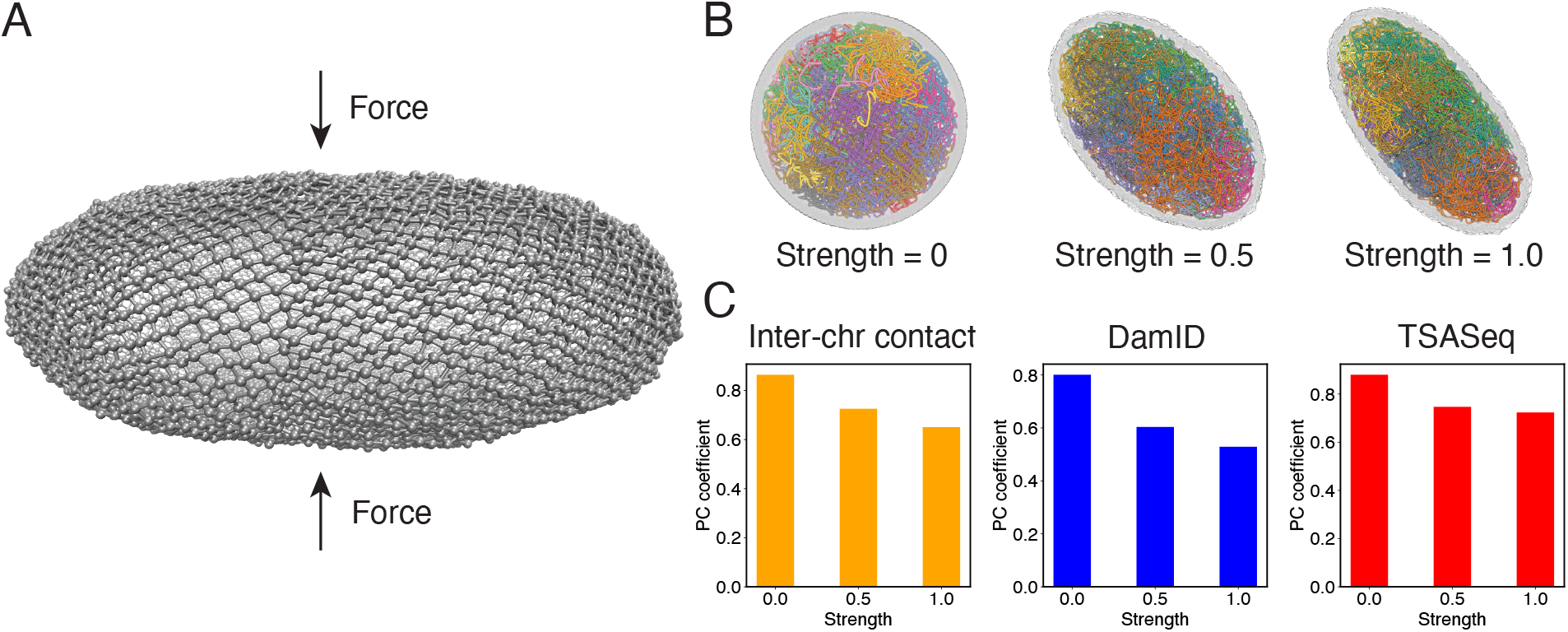
Nuclear deformations influence genome organization while preserving chromatin-speckle contacts. (A) Illustration of force induced nuclear envelope deformation. The nuclear lamina is modeled as a particle mesh where neighboring lamina particles are covalently bonded together. (B) Example nucleus conformations at different strengths of applied force. (C) Pearson correlation coefficients between results from simulations of deformed nuclei and those from a spherical nucleus for interchromosomal contacts (left), DamID profiles (middle), and TSA-Seq (right). The values at zero force were computed from two independent simulations starting from the same initial configurations.

As illustrated in Fig. 7B, the presence of external forces resulted in significant alterations in nuclear shape. We conducted two independent simulations with different force strengths, leading to varying degrees of deformation in the nuclear lamina. This deformation, in turn, caused a reorganization of chromosomes, affecting their normalized radial positions and pairwise contacts (see Fig. S4 and Fig. 7C). We observed that more deformed nuclei exhibited lower correlation coefficients for interchromosomal contacts compared to results obtained from simulations in a spherical nucleus. Similarly, the DamID profiles exhibited significant variations upon nucleus deformation, whereas TSA-Seq signals were much less affected and remained highly correlated with the results from the spherical nucleus simulations.

Therefore, it appears that speckles, and potentially other nuclear condensates, can dynamically reorganize in response to changes in chromosome conformations to maintain contacts with genomic loci. This robustness in nuclear body contacts may be essential for ensuring the robust functioning of the genome in a population of cells with significant variability in nuclear shape.

## Conclusions and Discussion

We introduced a computational model, OpenNucleome, to facilitate simulations for the human nucleus at high structural and temporal resolution. We conducted extensive crossvalidation with experimental data to support the biological relevance of simulated 3D structures. Implementing the model into the MD package, OpenMM enables GPU acceleration for long-timescale simulations. Tutorials in the format of Python Scripts with extensive documentation are provided to facilitate the adoption of the model by the community.

Our software enhances the capabilities of existing genome simulation tools. ^40,69,70^ Specifically, OpenNucleome aligns with the design principles of Open-MiChroM, ^70^ prioritizing open-source accessibility while expanding simulation capabilities to the entire nucleus. Similar to software from the Alber lab,^69^ OpenNucleome offers high-resolution genome organization that faithfully reproduces a diverse range of experimental data. Furthermore, beyond static structures, OpenNucleome facilitates dynamic simulations with explicit representations of various nuclear condensates, akin to the model developed by Fujishiro and Sasai ^40^.

A significant advantage of OpenNucleome lies in its predictive power for dynamical information. For example, the model succeeded in reproducing the subdiffusive behavior of telomeres. We further showed that the dynamics of individual chromosomes are slow and their radial positions do not relax over the time course of a cell cycle. This is consistent with previous theoretical estimations on chromosome dynamics ^92^ and recent observations of solid behavior of chromatin *in vivo*.^93^ Live cell experiments that directly track the positions of multiple chromosomes could further validate/falsify this prediction. We anticipate the model will greatly facilitate the investigation of the dynamics of genomic loci and nuclear bodies and the interpreting of live cell imaging results.

Slow chromosome dynamics and a lack of conformational relaxation naturally result in the heterogeneity of chromosome radial positions across individual cells. This heterogeneity raises doubts about the notion that chromosome radial positions provide robust and reliable mechanisms for gene regulation.^2,94–96^ Instead, our results support the nuclear zoning model for gene regulation,^36^ where specific loci function as “fixed points” anchored to certain nuclear bodies in all cells. This anchoring mechanism robustly creates the desired molecular environment surrounding these genomic segments. Unlike chromosome radial positions, contacts between genomic loci and speckles can be robustly established in individual cells, as shown in our simulations. It was achieved through a nucleation process that attracts speckle particles towards specific loci due to specific interactions. Nucleation occurs much more rapidly than chromosome rearrangement due to the smaller size of speckle particles. The coupled self-assembly mechanism for chromosomes and nuclear bodies can similarly facilitate the formation of other nuclear zones for different kinds of fixed points.

Despite the heterogeneity in chromosome positions and interchromosomal contacts, the ensemble of nuclear structures as a whole is not random and exhibits conserved features. For example, on average, certain chromosomes remain closer to the nuclear envelope than others (see Fig. 5B). Similarly, the average contact frequency between certain chromosome pairs is higher than others, though this trend can be frequently violated in individual cells. How such conserved features arise as cells exit from the mitotic phase remains unclear and would be interesting for further explorations.

## Methods

### Molecular dynamics simulation details

We used the software package OpenMM^71^ to perform MD simulations in reduced units at constant temperature (*T* = 1.0). Unless otherwise specified, we froze the lamina particles and only propagated the dynamics of chromatin, nucleoli, and speckles.

Two integration schemes were used with a time step of *dt* = 0.005 to efficiently generate structural ensembles and produce realistic dynamical information, respectively. For simulations used in parameter optimization and building structural ensembles, we employed the Langevin integrator with a damping coefficient of *γ^−^*^1^ = 10.0. In the case of MSD calculations shown in Fig. 6, we utilized Brownian dynamics with a damping coefficient of *γ^−^*^1^ = 0.01. The higher damping coefficient provides a better approximation to the viscous nucleus environment, while the smaller value in the Langevin integrator facilitates conformational sampling with faster diffusion rates.

We employed the semi-grand Monte Carlo technique^97^ to simulate chemical transitions between two types of speckle particles. At every 4000 simulation steps, we attempt a total of *N*_Sp_ chemical reactions that converts one type of speckle particles to the other type with a probability of 0.2. *N*_Sp_ corresponds to the total number of speckle particles, and the switching probability was chosen to be comparable to the experimental phosphorylation rate. More details on the speckle dynamics are provided in the Supporting Information *Section: Speckles as phase-separated droplets undergoing chemical modifications*.

When deforming the nuclear envelope, we unfroze the lamina particles and evolved them dynamically as the rest of the nucleus. Bonded interactions among lamina particles held the nuclear envelope together as a particle mesh. A harmonic force along the *z*-axis was introduced to compress the particle mesh. More details are provided in the Supporting Information *Section: Nuclear envelope deformation simulations*.

For simulations used to optimize parameters, a total of 50 independent 3-million-step-long trajectories were performed. Configurations were recorded at every 2000 simulation steps for analysis. The first 500,000 steps of each trajectory were discarded as equilibration. For production simulations, we performed 50 independent 10-million-step long trajectories starting from different initial configurations. Nuclear structures were again recorded at every 2000 steps to determine statistical averages presented in the paper. An additional 8 simulations of 30 million steps in length were performed to compute telomere MSDs.

We mapped the reduced units to real units with the conversion of length scale *σ* = 385 nm and the time scale in Brownian dynamics simulations *τ* = 0.65 *s*. These conversions were determined by as detailed in the Supporting Information *Section: Unit Conversion*.

### Experimental data processing and analysis

We obtained the in situ Hi-C data, SON TSA-seq data, and Lamin-B DamID data of HFF cell lines from the 4DN data portal. The intra and interchromosomal interactions were calculated at 100 KB resolution with VC SQRT normalization applied to the interaction matrices. Hi-C data extraction and normalization were performed using Juicer tools.^98^ We followed the same processing and normalization method described in Ref. 99 to analyze TSA-seq data. Two biological replicates of Lamin-B DamID data were merged and the normalized counts over Dam-only control were used for analysis. The SON TSA-Seq and Lamin-B DamID data were processed at the 25 KB resolution and the average values at the 100 KB resolution were used in Fig. 4 for model validation.

## Acknowledgement

This work was supported by the National Institutes of Health (Grant R35GM133580).

## Data and materials availability

Hi-C data (https://data.4dnucleome.org, accession number: 4DNFIB59T7NN). SON TSA-seq data (https://data.4dnucleome.org, accession number: pulldown data 4DNEX6U8TS3Y, control data 4DNEXI7XUWFK). Lam-inB DamID data (https://data.4dnucleome.org, accession number 4DNESXZ4FW4T). The software is available at: available at https://github.com/ZhangGroup-MITChemistry/OpenNucleome.

