## Supporting Information for "OpenNucleome for high resolution nuclear structural and dynamical modeling"

### Components of the whole nucleus model

As outlined in the main text, the whole nucleus model consists of chromosomes, nucleoli, speckles, and the nuclear lamina. Below, we provide details on the particle-based representations of the various components, totaling 70542 coarse-grained beads. Abbreviations are frequently used for clarity in notation, with N for nucleus, La for lamina, No for nucleoli, and Sp for speckles.

#### Chromosomes as beads on the string polymers

We explicitly modeled the 46 human chromosomes as beads-on-a-string polymers. Each coarse-grained bead represents a 100 KB genomic segment, totaling 60642 beads for the genome. We assigned each bead as either compartment type $A$, $B$, $C$, or $N$. The compartment assignments for types $A$ and $B$ were extracted from the Hi-C contact matrix for HFF cells [1] using the cooltools software [2], and compartment $C$ were identified as centromeric regions based on the DNA sequence. Compartment $N$ denotes genomic regions that cannot be assigned as $A$, $B$, or $C$ due to a lack of Hi-C data.

#### The nuclear lamina as a particle-based mesh

The nuclear envelope provides an enclosure to confine DNA and a repressive environment to organize chromatin with specific interactions [3]. To account for the role of the nuclear lamina while keeping our model simple, we approximate it with discrete particles uniformly placed on a sphere.

Following our previous work [4], we used the Fibonacci grid to initialize the lamina particles, which form a uniform and almost equidistant network of lamina particles on the surface of the nucleus [5,6]. The Cartesian coordinates associated with the $i^{th}$ lamina particles are defined as

$$\begin{aligned} x_i &= 2R_{\mathrm{N}} \times (1 - \frac{i}{N_{\mathrm{La}} - 1}) \\ y_i &= \sqrt{R_{\mathrm{N}}^2 - x^2} \times \cos[i\Phi] \\ z_i &= \sqrt{R_{\mathrm{N}}^2 - x^2} \times \sin[i\Phi] \end{aligned} \tag{S1}$$

where $N_{\mathrm{La}} = 8000$ represents the number of lamina particles, $i \in \{0, 1, \ldots, N_{\mathrm{La}} - 2, N_{\mathrm{La}} - 1\}$, and $\Phi = \pi \times (3 - \sqrt{5})$ is the golden angle. We set $R_{\mathrm{N}} = 5\mu m$ as the radius of the human foreskin fibroblasts (HFF) cell nucleus.

#### Nucleoli as phase-separated droplets

Nucleoli have been shown to behave as liquid droplets that form through phase separation [7–9]. We modeled the droplets with coarse-grained beads. While the composition of nucleoli is rather complex, we only used one type of particle for simplicity. In our simulations, we fixed the number of nucleolus particles, $N_{\mathrm{No}}$, based on the experimental concentration of nuclear protein NPM1, $c = 1\mu$M [4,10,11]. For example,

$$N_{\mathrm{No}} = \frac{4\pi}{3} \cdot N_{\mathrm{A}} \cdot (R_{\mathrm{No}})^3 \cdot c \approx 300 \tag{S2}$$

where $N_{\mathrm{A}}$ is the Avogadro constant and $R_{\mathrm{No}} = 0.5\ \mu m$ is the average nucleolous size [12,13].

##### Speckles as phase-separated droplets undergoing chemical modifications

Similar to nucleoli, speckles have also been shown to behave as liquid droplets [14]. However, one crucial unique feature of speckles is the constant chemical modifications of protein molecules comprising them, such as splicing factors [15]. The phosphorylation of these molecules has been argued to be essential for the dynamics and the number of speckles. Therefore, we implemented a kinetic scheme introduced by de Vries and coworkers [16] to account for the chemical reactions. In this scheme, we consider two types of speckle molecules: phosphorylated (Sp-P) and de-phosphorylated (Sp-dP). Only Sp-dP particles share attractive interactions.

The two protein types can inter-convert via chemical reactions with a transition probability matrix $\mathbf{T}$ defined as

$$\mathbf{T} = \begin{array}{r} \\ \text{Sp-dP} \\ \text{Sp-P} \end{array} \begin{array}{cc} \text{Sp-dP} & \text{Sp-P} \\ \left\| \begin{array}{cc} p_{11} & p_{12} \\ p_{21} & p_{22} \end{array} \right\| \end{array} = \begin{array}{r} \\ \text{Sp-dP} \\ \text{Sp-P} \end{array} \begin{array}{cc} \text{Sp-dP} & \text{Sp-P} \\ \left\| \begin{array}{cc} 0.8 & 0.2 \\ 0.2 & 0.8 \end{array} \right\| \end{array}. \tag{S3}$$

For simplicity, we assume the forward transition rate from Sp-P to Sp-dP particles is identical to the reverse rate. Because of the symmetry in transition rates, the average number of dP particles $\langle N_{\text{Sp-dP}} \rangle = 0.5 N_{\text{Sp}}$, where $N_{\text{Sp}}$ is the total number of speckle particles.

We chose the transition probability as 0.2 to be consistent with the phosphorylation rate. In particular, we estimate the rate as

$$k_{12} = p_{12} \times \tau^{-1} = 0.2 \times \frac{1}{4000 \times 0.005 \times 0.65s} = 0.0154\ s^{-1} \tag{S4}$$

where $\tau$ is the time interval between consecutive attempts of chemical reactions. As detailed in the *main text Section: Molecular dynamics simulation details*, the reactions were attempted every 4000 simulation steps, with a timestep of 0.005. The time unit in our simulations is 0.65s (see *Section: Mapping the Reduced Time Unit to Real Time*). The estimated value for $k_{12}$ is in the same order as the experimental phosphorylation rate [17].

We estimated the total number of speckle particles as follows. Assuming that there is a total of 30 speckles [18], we have $N_{\text{Sp-dP}} = 30 \times N_{\text{s}}$, where $N_{\text{s}}$ is the number of Sp-dP particles in each cluster. This estimation assumes that only Sp-dP particles share attractive interactions and contribute to cluster formation. From the experimentally estimated relative mass densities of the protein concentrations in the speckle and nucleolus droplet as $\frac{170}{203}$ [19], we have

$$\frac{N_{\text{s}} \times m/0.3^3}{100 \times m/0.5^3} = \frac{170}{203}. \tag{S5}$$

We assumed that speckle and nucleolus particles have identical mass and each nucleolus has 100 particles. The radius for speckle and nucleolus was approximated as 0.3 and $0.5\mu m$, yielding $N_s \approx 20$ and $N_{\text{Sp-dP}} \approx 600$. Because of the kinetic scheme defined in Eq. S3, only parts of Sp-dP particles will participate in droplet formation during the simulations. Therefore, we increase the particle number and set $\langle N_{\text{Sp-dP}} \rangle = 800$, which yields $N_{\text{Sp}} = 1600$.

#### Energy function of the whole nucleus model

As detailed below, the energy function of the whole nucleus, $U_{\text{Nucleus}}$, consists of interactions among chromosomes, among nuclear landmarks, and cross interactions between the two. Therefore,

$$U_{\text{Nucleus}} = U_{\text{Genome}} + U_{\text{NL}} + U_{\text{GN}} \tag{S6}$$

#### Hi-C inspired interactions for the diploid human genome

The energy function of the genome model is defined as

$$U_{\text{Genome}} = U_{\text{homo}}(\mathbf{r}) + U_{\text{ideal}}(\mathbf{r}) + U_{\text{compt}}(\mathbf{r}) + U_{\text{inter}}(\mathbf{r}). \tag{S7}$$

$U_{\text{homo}}(\mathbf{r})$ determines a generic polymeric topology of chromosomes with excluded volume effect:

$$U_{\text{homo}}(\mathbf{r}) = \sum_i \left[u_{\text{bond}}(r_{i,i+1}) + u_{\text{angle}}(\vec{r}_{i,i+1}, \vec{r}_{i+1,i+2})\right] + U_{\text{sc}}(\mathbf{r}) \tag{S8}$$

where the subscripts $i, i+1$, and $i+2$ represent the index of $i^{th}$, $(i+1)^{th}$, and $(i+2)^{th}$ beads, respectively, and $u_{\text{bond}}(r_{i,i+1})$ and $u_{\text{angle}}(r_{i,i+1}, r_{i+1,i+2})$ denote the bonding and angular potential applied for neighboring beads to ensure the connectivity of the chromatin chain and follow:

$$\begin{aligned} u_{\text{bond}}(r_{i,i+1}) &= K_2(r-r_0)^2 + K_3(r-r_0)^3 + K_4(r-r_0)^4, K_2 = K_3 = K_4 = 20\epsilon \\ u_{\text{angle}}(\vec{r}_{i,i+1}, \vec{r}_{i+1,i+2}) &= K_a\left[1-\cos(\theta-\pi)\right], K_a = 2\epsilon, \cos\theta = \frac{\vec{r}_{i,i+1}\cdot\vec{r}_{i+1,i+2}}{|\vec{r}_{i,i+1}|\cdot|\vec{r}_{i+1,i+2}|} \end{aligned} \tag{S9}$$

where, as discussed in Eq. S32, $r_0 = 0.5\sigma$ represents the size of the chromatin bead. The soft-core potential provides excluded volume effects for pairs of beads from the same or different chromosomes and follows:

$$U_{\text{sc}}(\mathbf{r}) = \sum_{j>i} u_{\text{sc}}(r_{ij}) \tag{S10}$$

where $u_{\text{sc}}(r_{ij})$ denotes a soft-core potential added to each pair formed by beads index $i$ and $j$ to account for the excluded volume effect while allowing the finite probability of cross-over of polymer chains.

$$u_{\text{sc}}(r_{ij}) = \begin{cases} 0.5E_{\text{cut}}\left(1+\tanh\left[\frac{2U_{\text{LJ}}(r_{ij})}{E_{\text{cut}}}-1\right]\right), & r_{ij}\le r_{\text{cut}} \\ U_{\text{LJ}}(r_{ij}), & r_{\text{cut}} < r_{ij} \le 0.5\times 2^{1/6}\sigma \\ 0, & r_{ij} > 0.5\times 2^{1/6}\sigma \end{cases} \tag{S11}$$

which corresponds to the Lennard-Jones potential capped off at a finite volume within a repulsive core to allow for chain crossing at a finite energy cost. $E_{\text{cut}} = 4\epsilon$ and $r_{\text{cut}}$ is chosen as the distance at which $U_{\text{LJ}}(r) = 0.5E_{\text{cut}}$.

$U_{\text{ideal}}(\mathbf{r})$ is the intra-chromosomal potential applied to genomic loci within the same chromosome, while $U_{\text{compt}}(\mathbf{r})$ is the compartment-specific interaction potential. The ideal potential, which can be rigorously derived following the maximum entropy principle [20, 21], adopts the following form:

$$U_{\text{ideal}}(\mathbf{r}) = \sum_I \sum_{i,j\in I} \alpha_{\text{ideal}}(|i-j|) f(r_{ij}) \tag{S12}$$

where $I$ indexes over each chromosome and $i$ and $j$ index over pair of beads on that chromosome. $\alpha_{\text{ideal}}(|i-j|)$ depends only on the sequence separation between two beads $i$ and $j$. $f(r_{ij})$ measures the probability of contact formation for two loci separated by a distance of $r_{ij}$, and its ensemble average corresponds to the contact probability measured in Hi-C experiments. $f(r_{ij})$ adopts the form:

$$\begin{aligned} f(r_{ij}) &= \frac{1}{2}\left(1+\tanh\left[(r_c-r_{i,j})^{-5} + 5(r_c-r_{i,j})\right]\right) \\ &+ \frac{1}{2}\left(1-\tanh\left[(r_c-r_{i,j})^{-5} + 5(r_c-r_{i,j})\right]\right)\times\left(\frac{r_c}{r_{i,j}}\right)^4. \end{aligned} \tag{S13}$$

The numerical value of $r_c$ was determined from the Hi-C contact map, as detailed in the next section. This contact probability function depicts that when $r < r_c, f \approx 1$ but when $r > r_c, f \approx (r_c/r)^4$. The power-law decay with an exponent of 4 is consistent with the relationship between contact probability and spatial distances revealed in imaging studies [22,23]. The tanh function ensures the continuity of the function and its derivative around $r_c$ (Fig. S5). Additionally, we truncated the ideal potential to be applicable for a sequence separation less than or equal to 100 MB and set the parameters for larger sequence separations to be zero. As shown in Fig. 4 of the main text, our parameterized ideal potential produced chromosomes with sizes comparable to imaging results. Incorporating longer-range interactions to improve the model further is straightforward but would also significantly increase the number of parameters.

Similar to the ideal potential discussed above, we have

$$U_{\text{compt}}(\mathbf{r}) = \sum_{i,j} \alpha_{\text{compt}}(T_i, T_j) f(r_{ij}), \tag{S14}$$

where $T_i$ and $T_j$ denote the compartment types of beads $i$ and $j$ which can be $A$, $B$ or $C$. Therefore, CG beads of the same compartment types will share the same interaction parameter $\alpha_{\text{compt}}(T_i, T_j)$, which will be derived from average Hi-C contact frequencies as detailed in the following sections.

To account for specific interactions between chromosomes, we introduced the interchromosomal potential as

$$U_{\text{inter}}(\mathbf{r}) = \sum_{I,J>I} \sum_{i\in I, j\in J} \alpha_{\text{inter}}(I, J) f(r_{ij}). \tag{S15}$$

$I, J \in \{1, 2, \ldots, 23\}$ index the haploid chromosomes, and parental and maternal chromosomes share identical parameters. This potential allows the model to capture interactions beyond those arising purely from compartmentalization as defined in Eq. S14.

All parameters in the energy function are summarized in Table S1. The procedure used for parameter optimization is detailed in the following sections.

#### Nuclear landmark-nuclear landmark interactions

The general energy function for interactions among nuclear landmark particles is defined as

$$U_{\text{NL}} = U_{\text{La}}(\mathbf{r}) + U_{\text{No}}(\mathbf{r}) + U_{\text{Sp-dP}}(\mathbf{r}) + U_{\text{EV}}(\mathbf{r}). \tag{S16}$$

The nuclear lamina was modeled as a particle mesh, and bonded potentials were introduced for nearest neighbor particles defined as

$$U_{\text{La}}(\mathbf{r}) = \sum_i \sum_{j\in\text{n.n.}, j>i} K_2(r - r_o)^2 + K_3(r - r_o)^3 + K_4(r - r_o)^4, K_2 = K_3 = K_4 = 20\epsilon \tag{S17}$$

with $r_o = 0.5\sigma$. $i$ indices all the lamina particles, and $j$ represents the nearest four neighbors around $i$ determined from the initial configuration for which the particles were placed on a Fibonacci grid. To avoid pairs $(i, j)$ being counted twice or more, we set $j$ always larger than $i$.

Short-ranged, non-bonded interactions were introduced among nuclear landmark particles to account for attractions that promote phase separation and the excluded volume effect. These interactions were modeled with a cut and shifted Lennard-Jones (LJ) potential defined as

$$U_{\text{LJ}}(r_{ij}) = \begin{cases} 4\epsilon_{\text{LJ}}\left(\left(\frac{\sigma_{\text{LJ}}}{r_{ij}}\right)^{12} - \left(\frac{\sigma_{\text{LJ}}}{r_{ij}}\right)^{6} - E_{\text{cut}}\right), & \text{for } r <= r_{\text{cut}} \\ 0, & \text{for } r > r_{\text{cut}} \end{cases} \tag{S18}$$

with $E_{\text{cut}} = 4\epsilon\left(\left(\frac{\sigma}{r_{\text{cut}}}\right)^{12} - \left(\frac{\sigma}{r_{\text{cut}}}\right)^{6}\right)$. We note that when $r_{\text{cut}}$ was set as $\sigma_{\text{LJ}} \times 2^{1/6}$, the potential has no attractive regime and only serves to prevent the overlap among particles, i.e., the excluded volume effect.

For attractive interactions between nucleolus particles, and between type dP speckle particles, we set the parameters as $\epsilon_{\text{LJ}} = 3.0$, $\sigma_{\text{LJ}} = 0.5$, and $r_{\text{cut}} = 1.5$. Therefore,

$$\begin{aligned} U_{\text{No}}(\mathbf{r}) = & \sum_{j>i\in\text{No}} U_{\text{LJ}}(r_{ij}, \epsilon_{\text{LJ}} = 3.0, \sigma_{\text{LJ}} = 0.5, r_{\text{cut}} = 1.5) \\ U_{\text{Sp-dP}}(\mathbf{r}) = & \sum_{j>i\in\text{Sp-dP}} U_{\text{LJ}}(r_{ij}, \epsilon_{\text{LJ}} = 3.0, \sigma_{\text{LJ}} = 0.5, r_{\text{cut}} = 1.5), \end{aligned} \tag{S19}$$

where the sums iterate over pairs of nucleolus particles and speckle dP particles.

For the excluded volume effect between nucleolus and speckle particles, between dP and P particles, and between P particles, we set the parameters as $\epsilon_{\text{LJ}} = 1.0$, $\sigma_{\text{LJ}} = 0.5$, and $r_{\text{cut}} = 0.5 \times 2^{1/6}$. These potentials are consistent with the estimated size of 0.5 $\sigma$ for speckle and nucleolus particles.

The excluded volume effect was also introduced between lamina and nucleolus particles and between the lamina and speckle particles to confine the nuclear bodies inside the nuclear envelope. We set the parameters as $\epsilon_{\text{LJ}} = 1.0$, $\sigma_{\text{LJ}} = 0.75$, and $r_{\text{cut}} = 0.75 \times 2^{1/6}$. The value for $\sigma_{\text{LJ}}$ was chosen based on a linear combination of the lamina particle size (1.0 $\sigma$) and the speckle/nucleolus particle size (0.5 $\sigma$).

Therefore, the excluded volume potential can be written as

$$\begin{aligned} U_{\text{EV}}(\mathbf{r}) = & \sum_{i\in\text{La}}\sum_{j\in\text{No}} U_{\text{LJ}}(r_{ij}, \epsilon_{\text{LJ}} = 1.0, \sigma_{\text{LJ}} = 0.75, r_{\text{cut}} = 0.75 \times 2^{1/6}) \\ + & \sum_{i\in\text{La}}\sum_{j\in\text{Sp}} U_{\text{LJ}}(r_{ij}, \epsilon_{\text{LJ}} = 1.0, \sigma_{\text{LJ}} = 0.75, r_{\text{cut}} = 0.75 \times 2^{1/6}) \\ + & \sum_{i\in\text{No}}\sum_{j\in\text{Sp}} U_{\text{LJ}}(r_{ij}, \epsilon_{\text{LJ}} = 1.0, \sigma_{\text{LJ}} = 0.5, r_{\text{cut}} = 0.5 \times 2^{1/6}) \\ + & \sum_{i\in\text{Sp-P}}\sum_{j\in\text{Sp-dP}} U_{\text{LJ}}(r_{ij}, \epsilon_{\text{LJ}} = 1.0, \sigma_{\text{LJ}} = 0.5, r_{\text{cut}} = 0.5 \times 2^{1/6}) \\ + & \sum_{j>i\in\text{Sp-dP}} U_{\text{LJ}}(r_{ij}, \epsilon_{\text{LJ}} = 1.0, \sigma_{\text{LJ}} = 0.5, r_{\text{cut}} = 0.5 \times 2^{1/6}) \\ + & \sum_{j>i\in\text{La}} U_{\text{LJ}}(r_{ij}, \epsilon_{\text{LJ}} = 1.0, \sigma_{\text{LJ}} = 0.5, r_{\text{cut}} = 0.5 \times 2^{1/6}). \end{aligned} \tag{S20}$$

We used abbreviations to denote various nuclear landmarks, with La for the nuclear lamina, Sp-P for P-type speckle particles, Sp-dP for dP-type speckle particles, and No for nucleolus particles. All the interaction parameters for the nuclear landmarks are listed in Table S3 for convenient reference.

#### Chromosome-nuclear landmark interactions

The energy function for interactions between chromosome and nuclear landmark particles is defined as

$$U_{\text{GN}} = U_{\text{C-La}}(\mathbf{r}) + U_{\text{C-Sp}}(\mathbf{r}) + U_{\text{C-No}}(\mathbf{r}). \tag{S21}$$

The functional form of the potential used to describe interactions between chromosomes and nuclear landmarks is inspired by experimental techniques that probe their contacts, such as lamin B DamID and SON TSA-Seq. For example, the average contact probability between a chromatin bead $i$ and the nuclear lamina can be estimated as

$$p_i^{\mathrm{L}} = \left\langle \sum_{j\in \mathrm{La}} c(r_{ij}) \right\rangle , \tag{S22}$$

where $j$ indexes over the lamina particles. $c(r_{ij})$ is defined as

$$c(r_{ij}) = \frac{1}{2}\Big(1 + \tanh\left[\eta(r_c - r_{ij})\right]\Big). \tag{S23}$$

It is a switching function that approaches one for $r_{ij} < r_c$, a threshold distance at which we set chromatin and the lamina as in contact. We chose $\eta = 4.0$ to obtain a reasonable decay of contact probability between chromosomes and nuclear landmarks. $r_c = 0.75$ was selected as the average size of the lamina (1.0 $\sigma$) and chromatin (0.5 $\sigma$) particles.

For the computational model to reproduce the experimental contact probability, following the maximum entropy argument [20, 21], the interaction potential between chromosomes and the nuclear lamina adopts the following form

$$\begin{aligned} U_{\text{C-La}}(\mathbf{r}) = \sum_{i\in \mathrm{Chr}} \sum_{j\in \mathrm{La}} \Big\{ &\frac{1}{2}\alpha_i^{\text{C-La}}\Big(1 + \tanh\left[\eta(r_c - r_{ij})\right]\Big) \\ &+ U_{\mathrm{LJ}}(r_{ij}, \epsilon_{\mathrm{LJ}} = 1.0, \sigma_{\mathrm{LJ}} = 0.75, r_{\mathrm{cut}} = 0.75 \times 2^{1/6})\Big\}. \end{aligned} \tag{S24}$$

A similar argument to the one outlined above was used to derive the interactions among chromosomes from Hi-C data, i.e., Eqs. S12, S14, and S15 [1]. The individual parameters $\alpha_i^{\text{C-La}}$ were optimized to ensure a match between simulated and experimental lamin B DamID data. The second term was included to account for the excluded volume effect and prevent chromatin from moving outside the envelope.

The interaction potential between chromosomes and the speckles adopts a similar form defined as

$$U_{\text{C-Sp}}(\mathbf{r}) = \sum_{i\in \mathrm{Chr}} \sum_{j\in \text{SP-dP}} \frac{1}{2}\alpha_i^{\text{C-Sp}}\Big(1 + \tanh\left[\eta(r_c - r_{ij})\right]\Big). \tag{S25}$$

The second sum for $j$ only includes dP-type speckle particles. The individual parameters $\alpha_i^{\text{C-Sp}}$ were optimized to ensure a match between simulated and experimental SON TSA-seq data.

Finally, the interaction potential between chromosomes and nucleoli is defined as

$$U_{\text{C-No}}(\mathbf{r}) = \sum_{i\in \mathrm{Chr}} \sum_{j\in \mathrm{No}} \frac{1}{2}\alpha_i^{\text{C-No}}\Big(1 + \tanh\left[\eta(r_c - r_{ij})\right]\Big). \tag{S26}$$

Because of the low data quality for the ChIP-Seq experiments for detecting chromatin-nucleoli contacts, we did not perform systematic optimizations for $\alpha_i^{\text{C-No}}$. Instead, we simply set them as $\alpha_i^{\text{C-No}} = P_i^{\mathrm{N}}\epsilon$, with $\epsilon = 1.0$. $P_i^{\mathrm{N}}$ is the probability for the chromatin bead $i$ to contact nucleoli as quantified by the software SPIN [24].

We list all the interaction parameters between chromosomes and the nuclear landmarks in Table S4.

#### Optimization of the whole nucleus model parameters

Below, we describe the procedures used to derive model parameters.

##### Connecting imaging and Hi-C data with the contact function

The function $f(r)$ defined in Eq. S13 was used to determine the chromatin contact probabilities. The availability of spatial positions and Hi-C data makes possible the definition of a contact function, $f(r)$, that converts distances into contact probabilities. In particular, we determined $r_c$ as the value at which the simulated average interchromosomal contact probability $\langle f(r_c)\rangle^{\text{sim}}_{\text{inter}}$ matches the experimental value, i.e.,

$$\langle f(r)\rangle^{\text{sim}}_{\text{inter}} = f^{\text{exp}}_{\text{inter}}. \tag{S27}$$

The angular brackets represent ensemble averaging, performed using the structures at 100 KB resolution reported in our previous work [4]. Matching simulation and experimental values produced $r_c = 0.54\sigma \approx 208$ nm. We note that this estimation for $r_c$ is comparable to the average bond length (0.5 $\sigma$), thus ensuring that nearest neighbor genomic regions with contact probability close to 1, i.e., $\langle f(r_{i,i+1})\rangle \approx 1$.

##### Adam optimizer for chromosome interaction parameters

Mathematical expressions for the various energy terms in $U_{\text{Genome}}$ were designed such that their ensemble averages can be mapped onto combinations of contact frequencies measured in Hi-C. The correspondence between the energy functions and Hi-C measurements allows model parameterization with an efficient adaptive moment (Adam) algorithm [25]. Specifically, $\alpha_{\text{ideal}}(|i-j|)$, $\alpha_{\text{compt}}(T_i, T_j)$, and $\alpha_{\text{inter}}(I, J)$ were tuned to satisfy the following constraints:

$$\begin{aligned}
\left\langle \sum_I \sum_{i,j\in I} f(r_{ij})\delta_{|i-j|,s} \right\rangle &= \sum_I \sum_{i,j\in I} f^{\text{exp}}_{ij}\delta_{|i-j|,s}, \qquad \text{for } s = 1,\cdots,n-1 \\
\left\langle \sum_{i,j} f(r_{ij})\delta_{T_i,T_1}\delta_{T_j,T_2} \right\rangle &= \sum_{i,j} f^{\text{exp}}_{ij}\delta_{T_i,T_1}\delta_{T_j,T_2}, \qquad \text{for } T_1, T_2 \in \{A, B, C\} \\
\left\langle \sum_{i\in I, j\in J} f(r_{ij})\delta_{I,K_1}\delta_{J,K_2} \right\rangle &= \sum_{i\in I, j\in J} f^{\text{exp}}_{ij}\delta_{I,K_1}\delta_{J,K_2}, \\
&\qquad\qquad \text{for } K_1, K_2 \in \{1, \ldots, 23\}
\end{aligned} \tag{S28}$$

where $\delta_{T_i,T_1}$ is the Kronecker delta function with the following definition:

$$\delta_{T_i,T_1} = \begin{cases} 1, & \text{if } T_i = T_1 \\ 0, & \text{otherwise} \end{cases} \tag{S29}$$

The angular bracket represents the ensemble average, and $f^{\text{exp}}_{ij}$ is the corresponding experimental contact frequency.

During the optimization process, our aim was to minimize the disparity between experimental findings and simulated data. To achieve this, we defined the cost function as follows:

$$L = \sum_i (\langle f_i\rangle - f^{\text{exp}}_i)^2, \tag{S30}$$

where the index $i$ iterates over all the constraints defined in Eq. S28.

The details of the algorithm for parameter optimization are as follows:

1. Starting with a set of values for $\alpha_{\text{ideal}}(|i-j|)$, $\alpha_{\text{compt}}(T_i, T_j)$, and $\alpha_{\text{inter}}(I, J)$, we performed 50 independent 3-million-step long MD simulations to obtain an ensemble of nuclear configurations. The 500K steps of each trajectory are discarded as equilibration. We collected the configurations at every 2000 simulation steps from the rest of the simulation trajectories to compute the ensemble averages defined on the left-hand side of Eq. S13.

2. Check the convergence of the optimization by calculating the percentage of error defined as $\sum_i (\langle f_i \rangle - f_i^{\text{exp}}) / \sum_i f_i^{\text{exp}}$. The summation over $i$ includes all the average contact probabilities defined in Eq. S28.

3. If the error is less than a tolerance value $e_{\text{tol}}$, the optimization has converged, and we stop the simulations. Otherwise, we update the parameters, $\alpha$, using the Adam optimizer [25]. With the new parameter values, we return to step one and restart the iteration.

#### Adam optimizer for chromosome-nuclear body interaction parameters

Similar to those among chromatin particles, the interaction parameters between chromatin and nuclear landmarks were optimized with Adam's algorithm to reproduce experimental constraints.

The constraints that we aimed to reproduce were defined as follows,

$$\begin{aligned} \left\langle C_i^{\text{La}} \right\rangle &= \text{LAF}_i^{\text{exp}}, \quad \text{for } i = 1, \cdots, N \\ \left\langle C_i^{\text{Sp}} \right\rangle &= \text{SAF}_i^{\text{exp}}, \quad \text{for } i = 1, \cdots, N, \end{aligned} \tag{S31}$$

where $C_i^{\text{La}}$ and $C_i^{\text{Sp}}$ measure the contacts between chromatin bead $i$ and nuclear lamina and speckles, respectively, as defined in Eq. S42 and S45. $\text{LAF}_i$ and $\text{LAF}_i$denote the lamina and speckle association frequency for chromatin bead $i$ as measured in Lamin B DamID and SON TSA-Seq experiments. $N$ denotes the number of chromatin beads. We combined the constraints defined in Eq. S31 with those in Eq. S28 to simultaneously optimize the parameters using the iterative algorithm outlined in the previous section. We note that the interaction potential between chromatin and speckles defined in Eq. S25 did not use precisely the same function as in $C_i^{\text{Sp}}$. We chose to sum over all speckle dP particles, rather than identifying the droplets, which is difficult to do during the simulations.

#### Parameter optimization for nuclear body-nuclear body interactions

As much remains to be known about the organization of nuclear bodies, we designed the interaction potentials and parameters based on qualitative observations without extensive fine-tuning. For example, we used the standard Lennard-Jones potential (Eq. S18) to mimic short-range interactions. The lengthscales, $\sigma_{\text{LJ}}$, in these potentials, were chosen based on a linear combination of the size of interacting particles, as discussed in *Section: Unit Conversion*.

The interaction strength, $\epsilon_{\text{LJ}}$, was set as 1.0 to be on the same order as thermal energy ($k_{\text{B}}T$), when the potential was used to account for the excluded volume effect.

For attractive interactions that promote phase separation and nuclear body formation, we set $\epsilon_{\text{LJ}} = 3.0$. Smaller values failed to produce clustered nucleoli, while much larger values significantly decreased the fluidity of the resulting droplets. The same value was used for speckle dP particles and produced droplet numbers comparable to experimental observations (Fig. S1).

#### Unit Conversion

The reduced unit for length scale is noted as $\sigma$. We set the nucleus radii as $13\sigma$. Assuming a nucleus with an average size of 5 $\mu$m, we have $\sigma = 385$nm.

##### Mapping chromatin bead size to real unit

We estimated the size of the chromosome bead as 192.5 nm based on super-resolution imaging data as follows. The median radius of gyration has been shown to follow a power-law scaling as a function of domain length with an exponent of 0.3 [26]. Assuming that the radius of a domain is proportional to the radius of gyration, we have

$$R \propto R_g \propto L^{0.3} \Longrightarrow \frac{R_{\mathrm{1MB}}}{R_{100KB}} = \Big(\frac{1MB}{100KB}\Big)^{0.3}. \tag{S32}$$

We previously estimated the size of 1MB bead as $R_{\mathrm{1MB}} = \sigma = 385$nm, and Eq. S32 yields the size of 100 KB as $R_{\mathrm{100KB}} = 0.5\sigma$.

##### Mapping lamina bead size to real unit

We chose the number and the diameter of lamina beads $N_{\mathrm{La}}, \sigma_{\mathrm{La}}$ by estimating the distance between nearest neighbor lamina beads. We found that at $N_{\mathrm{La}} = 8000$, when the lamina particles were placed on the Fibonacci grid over the spherical surface, the average nearest neighbor distance was 0.52. Therefore, we set $\sigma_{\mathrm{La}} = 0.5\sigma$ when considering the excluded volume effect between lamina particles. However, when modeling the excluded volume effect between lamina and chromatin, nucleolus, or speckle particles, we used $\sigma_{\mathrm{La}} = 1.0$ (see Eq. S20). A larger value provides a stronger excluded volume effect that prevents these particles from crossing the nucleus boundary or getting stuck in the space of the lamina particle mesh grid.

##### Mapping nucleoli bead size to real unit

The size of nucleolus particles ($\sigma_{\mathrm{No}}$) was estimated as follows. Since the average number of nucleoli inside a cell nucleus ranges from 2-5, we approximate the number of particles comprising individual droplets as $N_{\mathrm{No}}/3$, assuming a total of three nucleoli. $N_{\mathrm{No}}$ corresponds to the total number of nucleolus particles. With a space-filling model, the ratio of the volume between one nucleolus and the cell nucleus can be estimated as

$$\frac{(4\pi/3)(2^{1/6}\sigma_n/2)^3(N_{\mathrm{No}}/3)}{(4\pi/3)R_{\mathrm{N}}^3} = \Big(\frac{R_{\mathrm{No}}}{R_{\mathrm{N}}}\Big)^3 \tag{S33}$$

where $2^{1/6}\sigma_n/2$ denotes the effective radius of a nucleolus particle, and $R_{\mathrm{N}}$ is the nucleus size. Using experimental values for the nucleolus and nucleus size [12,13] as $R_{\mathrm{No}} = 0.5\mu m$ and $R_{\mathrm{N}} = 5\mu m$, we have $\sigma_{\mathrm{No}} = 0.5$.

##### Mapping speckle bead size to real unit

A similar procedure as in the previous section was used to estimate the size of speckle particles $\sigma_{\mathrm{Sp}}$. Since approximately 600 dP-type speckle particles form speckle clusters, each speckle cluster consists of around 20 particles. This estimation assumes a total of 30 speckle droplets in the system, consistent with the experimentally reported range of 20-50 speckles. [18] With a space-filling model, the ratio of the volume between one speckle and the cell nucleus can be estimated as

$$\frac{(4\pi/3)(2^{1/6}\sigma_{\mathrm{Sp}}/2)^3(N_{\mathrm{Sp}})}{(4\pi/3)R_{\mathrm{N}}^3}=\Big(\frac{R_{\mathrm{Sp}}}{R_{\mathrm{N}}}\Big)^3 \tag{S34}$$

where $N_{\mathrm{Sp}}=20$. Using experimental values for the speckle and nucleus size [19] as $R_{\mathrm{Sp}}=0.3\mu m$ and $R_{\mathrm{N}}=5\mu m$, we have $\sigma_{\mathrm{Sp}}=0.5$.

##### Mapping the reduced time unit to real time

We determined the timescale mapping by matching the simulated diffusion coefficient of chromatin particles with experimental values. The diffusion coefficient in our simulations can be estimated from the fluctuation-dissipation theorem [27] as $D=\frac{k_BT}{\zeta}$, where the friction coefficient $\zeta=m\gamma$. Using the conversion from $\frac{k_{\mathrm{B}}T}{m}=\frac{\sigma^2}{\tau_{\mathrm{B}}^2}$, we have

$$D=\frac{k_BT}{\zeta}=\frac{k_BT}{m\gamma}=\frac{\sigma^2}{\tau_{\mathrm{B}}^2\gamma}=\frac{10^{-2}\sigma^2}{\tau_B}. \tag{S35}$$

We used the simulation setup $\gamma^{-1}=10^{-2}\tau_B$ when deriving the last equation.

In the meantime, from the Stokes-Einstein (SE) equation, we have $D=\frac{k_BT}{6\pi\eta r}$, where $\eta$ is the viscosity and $r=0.25\sigma$ is the radius of chromatin beads. Therefore,

$$\frac{k_BT}{6\pi\eta r}=\frac{10^{-2}\sigma^2}{\tau_B}, \tag{S36}$$

and

$$\tau_B=\frac{10^{-2}\sigma^2\cdot 6\pi\eta r}{k_BT}=\frac{1.5\times10^{-2}\pi\eta\sigma^3}{k_BT}. \tag{S37}$$

Setting the nucleoplasmic viscosity as $1Pa\cdot s$ produces $\tau_B\approx 0.65s$. This mapping produced diffusion coefficients and mean squared displacement curves that match well with experimental measurements presented in Ref. [28], as discussed in the main text. We note that the chosen value for the nucleoplasmic viscosity indeed falls into the range of reported experimental values from $10^{-1}Pa\cdot s$ to $10^2Pa\cdot s$ [29,30].

#### Molecular dynamics simulation details

##### Initial configurations for simulations

Due to the slow relaxation dynamics of whole chromosomes relative to the simulation timescale, the reported results are sensitive to the configurations used to initialize the simulations. Therefore, we designed the following protocol to prepare the initial configurations and ensure the biological relevance of simulation results.

We first created a total of 1000 configurations for the genome by sequentially generating the conformation of each one of the 46 chromosomes as follows. For a given chromosome, we start by placing the first bead at the center (origin) of the nucleus. The positions of the following beads, $i$, were determined from the $(i-1)$-th bead as $\boldsymbol{r}_i=\boldsymbol{r}_{i-1}+0.5\boldsymbol{v}$. $\boldsymbol{v}$ is a normalized random vector, and 0.5 was selected as the bond length between neighboring beads. To produce globular chromosome conformations, we rejected vectors, $\boldsymbol{v}$, that led to bead positions with distance from the center larger than $4\sigma$. Upon creating the conformation of a chromosome $i$, we shift its center of mass to a value $r^i_{\text{com}}$ determined as follows. We first compute a mean radial distance, $r^i_{\text{o}}$ with the following equation

$$\frac{6\sigma - r^i_{\text{o}}}{r^i_{\text{o}} - 2\sigma} = \frac{D_{\text{hi}} - D}{D - D_{\text{lo}}}, \tag{S38}$$

where $D_i$ is the average value of Lamin B DamID profile for chromosome $i$. $D_{\text{hi}}$ and $D_{\text{lo}}$ represent the highest and lowest average DamID values of all chromosomes, and $6\sigma$ and $2\sigma$ represent the upper and lower bound in radial positions for chromosomes. As shown in Fig. S6, the average Lamin B DamID profiles are highly correlated with normalized chromosome radial positions as reported by DNA MERFISH [31], supporting their use as a proxy for estimating normalized chromosome radial positions. We then select $r^i_{\text{com}}$ as a uniformly distributed random variable within the range $[r^i_{\text{o}} - 2\sigma, r^i_{\text{o}} + 2\sigma]$. Without loss of generality, we randomly chose the directions for shifting all 46 chromosomes.

We further relaxed the 1000 configurations to build more realistic genome structures. Following an energy minimization process, one-million-step molecular dynamics (MD) simulations were performed starting from each configuration. Simulations were performed with the following energy function

$$U_{\text{Relax}} = U_{\text{Genome}} + U^{\text{EV}}_{\text{C-La}}, \tag{S39}$$

where $U_{\text{Genome}}$ is defined as in Eq. S7. $U_{\text{G-La}}$ is the excluded volume potential between chromosomes and lamina, i.e, only the second term in Eq. S24. Parameters in $U_{\text{Genome}}$ were from a preliminary optimization. The end configurations of the MD simulations were collected to build the final configuration ensemble (FCE).

We further computed the Pearson correlation coefficient of pairwise interchromosomal contacts between different structures in FCE (see *Section: Computing pairwise interchromosomal contact probabilities*). As shown in Fig. S3A, the probability distribution of these correlation coefficients is comparable with that determined from DNA-MERFISH structures, supporting the biological relevance of the structural diversity in the constructed ensemble.

From 1000 relaxed configurations, we selected a subset of structures to initialize simulations presented in the main text. An optimization procedure was introduced for structure selection. We start this procedure by randomly select $N$ structures to build the initial configuration ensemble (ICE). We then iteratively go through every configuration in ICE and replace with a structure from FCE that's not already included in ICE. We then compute the Pearson correlation coefficient between new average ICE interchromosomal contact probabilities and experimental values. If the Pearson correlation coefficient is higher than the value determined from the original ICE, the new structure is accepted and the ICE is updated. Otherwise, the new structure is rejected. We stop the selection process for when the Pearson correlation coefficient stops improving.

We found that as $N$ increases, the agreement between ICE interchromosomal contact probabilities and experimental values continue to increase (Fig. S3B). We set $N = 50$, which produces a Pearson correlation coefficient between ICE and experimental interchromosomal contact probabilities of 0.9. Further increasing $N$ does not significantly improve the agreement but incurs more computational cost.

It is worth noting that outcomes of the selection procedure depend on the initial set of configurations included in ICE at the beginning. However, we found that the ICEs produced from 20 independent trials are highly correlated (Fig. S3C) and all reproduce the heterogeneity in interchromosomal contacts seen in DNA MERFISH data (Fig. S3D). Therefore, the selection procedure is robust and can produce biologically meaningful configurations to initialize simulations.

With the chromosome positions prepared, we randomly placed 300 nucleoli and 1600 speckle particles inside the nucleus to complete the set up of initial configurations.

##### Langevin dynamics simulations

We used the Langevin integrator with the damping coefficient $\gamma^{-1} = 10$ to control the temperature at $T = 1.0$ for simulations used for parameter optimization and for producing an ensemble of nucleus structures. Langevin dynamics simulations allow faster chromosome movements, compared to Brownian dynamics simulations, facilitating the conformational sampling. In these simulations, the lamina particles were frozen and no explicit dynamics were considered for the nuclear envelope.

##### Brownian dynamics simulations

We also performed Brownian dynamics simulations with damping coefficient $\gamma^{-1} = 10^{-2}$ to control the temperature at $T = 1.0$. These simulations provide better approximations of the overdamped dynamics of chromatin for direct comparison with live cell imaging studies. As detailed in the *Section: Unit Conversion*, upon mapping the coarse-grained timescale to the physical unit, Brownian dynamics simulations produce diffusion coefficients for telomeres comparable to experimental values (see Fig. 5 of the main text).

##### Nuclear envelope deformation simulations

We performed Langevin dynamics simulations to investigate the impact of nuclear envelope deformation on genome organization. To induce a compressing force along the $z$-axis, we introduced a harmonic potential in the form of

$$U_{\text{compress}} = \sum_{i=1}^{N_{\text{La}}} k \times \frac{z_i^2}{R_N}. \tag{S40}$$

$z_i$ is the $z$ coordinate of the $i$-th lamina bead, and $N_{\text{La}}$ represents the total number of lamina beads. The particles in the system evolve under the combined effect of $U_{\text{compress}}$ and $U_{\text{Nucleus}}$ defined in Eq. S6.

#### Details of simulation data analysis

The computer simulations yield 3D coordinates of the diploid genome. However, when comparing directly with experimental data processed for the haploid genome, unless stated otherwise, we computed averages across paternal and maternal chromosomes to ascertain various genome-wide properties as listed below.

##### Computing simulated contact probabilities

Simulated contact probability maps were computed by averaging over chromosome configurations collected from all trajectories. For a given configuration, the contact probability between two chromatin segments ($i$ and $j$) was evaluated using the contact function defined in Eq. S13.

#### Computing the Pearson correlation coefficients between experimental and simulated contact maps

We computed the Pearson correlation coefficients (PCC) between experimental and simulated contact maps in Fig. 4A and Fig. S2 as

$$r = \frac{\sum_{i=1}^{n}(x_i - \bar{x})(y_i - \bar{y})}{\sqrt{\sum_{i=1}^{n}(x_i - \bar{x})^2}\sqrt{\sum_{i=1}^{n}(y_i - \bar{y})^2}}. \tag{S41}$$

$x_i$ and $y_i$ represent the experimental and simulated contact probabilities, and $n$ is the total number of data points. Only non-redundant data points, i.e., half of the pairwise contacts, are used in the PCC calculation.

#### Computing pairwise interchromosomal contact probabilities

For a given genome structure, we computed the pairwise interchromosomal contacts as follows. For every pair of chromosomes, we determined their contact probability by averaging all genomic pairs from two chromosomes using Eq. S13. We then averaged over all four pairs of diploid chromosomes to compute the haploid average contacts. In total, there are $C_{22}^2 = 231$ contact pairs between haploid chromosomes excluding the sex chromosomes.

#### Distances from nuclear bodies and association frequencies

The contacts of a chromatin bead $i$ with the nuclear lamina were evaluated as

$$C_i^{\text{La}} = \frac{1}{N_t}\sum_t \sum_{j\in\text{La}} \frac{1}{2}\Big(1 + \tanh[\eta(r_c - r_{i,j})]\Big) \tag{S42}$$

with $r_c = 0.75\sigma$. We average over the ensemble of nuclear configurations and homologs to compute the *in silico* Lamin B DamID signal as

$$\text{DamID}_i = \log_2\Big(\frac{\langle C_i^{\text{La}}\rangle}{\overline{C}^{\text{La}}}\Big), \tag{S43}$$

where the angular brackets indicate ensemble averaging. $\overline{C}^{\text{La}}$ is defined as the genome wide average of $\langle C_i^{\text{La}}\rangle$.

For chromatin-speckle contacts, we first identified the speckles formed at any given structure using the density-based spatial clustering algorithm DBSCAN [32] as implemented in the scikit library for Python [33]. For the identified droplets, we computed their center of mass coordinates, $\vec{r}^{\ com}$ and the radius of gyration, $R$. With the identified clusters, we then determined the distance from the $i$-th chromatin bead to the $s$-th speckle as

$$d_{i,s} = \|\vec{r}_i - \vec{r}_s^{\ \text{com}}\| - R_s, \tag{S44}$$

where $\|\cdot\|$ represents the L2 norm. We subtract the radius of the speckle cluster in the above equation to determine the distance to the droplet surface. From the list of distances to different speckles, the contact between chromatin bead $i$ and speckles is computed as

$$C_i^{\text{Sp}} = \frac{1}{N_s}\sum_s \frac{1}{2}\Big(1 + \tanh[\eta(d_c - d_{i,s})]\Big), \tag{S45}$$

where we sum over all the $N_s$ speckle clusters. A similar expression was used for determining the contacts between chromatin and nucleoli.

Finally, we average over the ensemble of nuclear configurations and homologs to compute the *in silico* SON TSA-Seq signal as

$$\mathrm{TSA_i} = \log_2\Big(\frac{\left\langle \mathrm{C_i^{Sp}} \right\rangle}{\overline{\mathrm{C}}^{\mathrm{Sp}}}\Big), \tag{S46}$$

where the angular brackets indicate ensemble averaging. $\overline{C}^{\mathrm{Sp}}$ is defined as the genome wide average of $\left\langle C_i^{\mathrm{Sp}} \right\rangle$.

##### Computing simulated normalized chromosome radial positions

For a given chromosome $i$, we first determined its center of mass position denoted as $C_i$. Starting from the center of the nucleus, $O$, we extend the the vector $\boldsymbol{v}_{OC}$ to identify the intersection point with the nuclear lamina as $P_i$. The normalized radial position of chromosome $i$ is then defined as $\frac{\|\boldsymbol{v}_{OC_i}\|}{\|\boldsymbol{v}_{OP_i}\|}$, where $\|.\|$ represents the L2 norm.

##### Computing simulated chromosome radii of gyration

The radius of gyration for a chromosome is computed as,

$$R_g = \sqrt{\frac{\sum_i^n \|\boldsymbol{r}_i - \boldsymbol{r}_{\mathrm{com}}\|}{n}}, \tag{S47}$$

where $\boldsymbol{r}_{\mathrm{com}}$ and $n$ are the center of mass and the number of beads of the chromosome. $i$ indices over all the chromosome beads and $\boldsymbol{r}_i$ correspond to the Cartesian coordinates of bead $i$. $\|.\|$ represents the L2 norm.

##### Computing simulated mean square displacement

Mean square displacements (MSD) for telomeres were computed as

$$\left\langle \boldsymbol{r}^2(\Delta t) \right\rangle = \frac{1}{N_{\mathrm{traj}}} \sum_{t=1}^{N_{\mathrm{traj}}} \frac{1}{N_{\mathrm{step}}} \sum_{i=1}^{N_{\mathrm{step}}} [\boldsymbol{r}^t((i-1)\delta t + \Delta t) - \boldsymbol{r}^t((i-1)\delta t)]^2, \tag{S48}$$

where $\Delta t, \delta t,$ and $N_{\mathrm{step}}$ represent the time interval, the time step, and the total number of steps, respectively. The summation over $t$ corresponds to averaging over eight independent trajectories. MSDs telomeres from paternal and maternal chromosomes are separately computed and analyzed.

#### Details of experimental data analysis

##### Interchromosomal contacts from DNA MERFISH data

We collected the DNA MERFISH data reported in Ref. [31] to construct the experimental ensemble of 5455 genome structures. For each structure, we computed the pairwise interchromosomal contacts following the procedure outlined in *Section: Computing pairwise interchromosomal contact probabilities.*

To better visualize and analyze interchromosomal contacts, we applied the Uniform Manifold Approximation and Projection (UMAP) technique as implemented in software package umap-learn [34, 35], with default parameters to reduce the 231 haploid contacts into two dimensions. All 5455 DNA MERFISH structures were included in this analysis.

The same transformations produced from the UMAP analysis of experimental structures were applied to in silico configurations to produce results shown in Fig. S3C and Fig. S3D.

#### Computing experimental normalized chromosome radial positions

We followed the same procedure outlined in *Section: Computing simulated normalized chromosome radial positions* to compute the experimental values. To determine the center of the nucleus using DNA MERFISH data, we used the algorithm, minimum volume enclosing ellipsoid (MVEE) [36], to fit an ellipsoid for each genome structure. The optimal ellipsoid defined as $(\boldsymbol{x}-\boldsymbol{c})^T\boldsymbol{A}(\boldsymbol{x}-\boldsymbol{c})\equiv 1$ is obtained by optimizing $\min(\log(\det[\boldsymbol{A}]))$ subjecting to the constraint that $(\boldsymbol{x}_i-\boldsymbol{c})^T A(\boldsymbol{x}_i-\boldsymbol{c})\leq 1$. $\boldsymbol{x}_i$ correspond to the list of chromatin positions determined experimentally.

#### Computing experimental radii of gyration

We computed the experimental radii of gyration with using the same expression as that for analyzing simulated structures (Eq. S47).

**Table S1. Summary of the various terms of the chromosome energy function and the algorithms used for parameter optimization.** See also *Section: Hi-C inspired interactions for the diploid human genome* for detailed expression of the energy function and *Section: Adam optimizer for chromosome interaction parameters* for details on the optimization algorithm.

| Potentials | Functional forms | Parameter values |
|---|---|---|
| Bonding potential | $u_{\text{bond}}(r_{i,i+1})$ in Eq. S9 | Standard values in coarse-grained polymer models |
| Angular Potential | $u_{\text{angle}}(\vec{r}_{i,i+1}, \vec{r}_{i+1,i+2})$ in Eq. S9 | Standard values in coarse-grained polymer models |
| Soft-core potential | $u_{\text{sc}}(r_{ij})$ in Eq. S11 | Standard values in coarse-grained polymer models |
| Ideal potential | $U_{\text{ideal}}(\mathbf{r})$ in Eq. S12 | Values for $\alpha_{\text{ideal}}$ were obtained from optimizations against Hi-C data (see Fig. S7). |
| Compartment potential | $U_{\text{compt}}(\mathbf{r})$ in Eq. S14 | Values for $\alpha_{\text{compt}}$ were obtained from optimizations against Hi-C data (see Table 2). |
| Inter potential | $U_{\text{inter}}(\mathbf{r})$ in Eq. S15 | Values for $\alpha_{\text{inter}}$ were obtained from optimizations against Hi-C data (see Fig. S8). |

**Table S2.** Summary of interaction parameters between various compartment types, i.e. $\alpha_{\text{compt}}$ defined in Eq. S14.

| | |
|---|---|
| $\alpha_{AA}$ | -0.074185 |
| $\alpha_{AB}$ | 0.112285 |
| $\alpha_{AC}$ | 0.009947 |
| $\alpha_{BB}$ | 0.059981 |
| $\alpha_{BC}$ | 0.072481 |
| $\alpha_{CC}$ | 0.088825 |

**Table S3. Summary of the interaction potentials among particles that make up the nuclear landmarks and their corresponding parameter values.** See also text *Section: Nuclear landmark-nuclear landmark interactions* for further discussion and *Section: Unit Conversion* for choosing the size of various particles, from which the $\sigma_{\text{LJ}}$ were derived with a linear combination rule.

| Potentials | Function forms | Parameter values |
|---|---|---|
| Nucleolus/ Nucleolus | $U_{\text{LJ}}(r_{ij})$ in Eq. S18 | $\epsilon_{\text{LJ}} = 3.0$, and $r_{\text{cut}} = 1.5$ were chosen to mimic short-range attractions that produce an average of two nucleoli per cell. $\sigma_{\text{LJ}} = \sigma_{\text{No}} = 0.5$. |
| Sp-dP/ Sp-dP (speckles) | $U_{\text{LJ}}(r_{ij})$ in Eq. S18 | $\epsilon_{\text{LJ}} = 3.0$ and $r_{\text{cut}} = 1.5$ were chosen to mimic short-range attractions that produce around 30 speckle droplets. $\sigma_{\text{LJ}} = \sigma_{\text{Sp-dP}} = 0.5$. |
| Sp-dP/ Sp-P (speckles) | $U_{\text{LJ}}(r_{ij})$ in Eq. S18 | $\epsilon_{\text{LJ}} = 1.0$ and $r_{\text{cut}} = 0.5 \times 2^{1/6}$ were chosen as standard values to provide the excluded volume effect. $\sigma_{\text{LJ}} = \frac{\sigma_{\text{Sp-dP}}+\sigma_{\text{Sp-P}}}{2} = 0.5$. |
| Sp-P/ Sp-P (speckles) | $U_{\text{LJ}}(r_{ij})$ in Eq. S18 | $\epsilon_{\text{LJ}} = 1.0$ and $r_{\text{cut}} = 0.5 \times 2^{1/6}$ were chosen as standard values to provide the excluded volume effect. $\sigma_{\text{LJ}} = \sigma_{\text{Sp-P}} = 0.5$. |
| Nucleolus/ Speckle | $U_{\text{LJ}}(r_{ij})$ in Eq. S18 | $\epsilon_{\text{LJ}} = 1.0$ and $r_{\text{cut}} = 0.5 \times 2^{1/6}$ were chosen as standard values to provide the excluded volume effect. $\sigma_{\text{LJ}} = \frac{\sigma_{\text{No}}+\sigma_{\text{Sp}}}{2} = 0.5$. |
| Nucleolus/ Lamina | $U_{\text{LJ}}(r_{ij})$ in Eq. S18 | $\epsilon_{\text{LJ}} = 1.0$ and $r_{\text{cut}} = 0.75 \times 2^{1/6}$ were chosen as standard values to provide the excluded volume effect. $\sigma_{\text{LJ}} = \frac{\sigma_{\text{No}}+\sigma_{\text{La}}^{\dagger}}{2} = 0.75$. |
| Speckle/ Lamina | $U_{\text{LJ}}(r_{ij})$ in Eq. S18 | $\epsilon_{\text{LJ}} = 1.0$ and $r_{\text{cut}} = 0.75 \times 2^{1/6}$ were chosen as standard values to provide the excluded volume effect. $\sigma_{\text{LJ}} = \frac{\sigma_{\text{Sp}}+\sigma_{\text{La}}^{\dagger}}{2} = 0.75$. |
| Lamina/ Lamina | $U_{\text{LJ}}(r_{ij})$ in Eq. S18 | $\epsilon_{\text{LJ}} = 1.0$ and $r_{\text{cut}} = 0.5 \times 2^{1/6}$ were chosen as standard values to provide the excluded volume effect. $\sigma_{\text{LJ}} = \sigma_{\text{La}} = 0.5$. |

$^{\dagger}$As mentioned in the *Section: Mapping lamina bead size to real unit*, a larger value for $\sigma_{\text{La}}$ was used here to provide a stronger excluded volume effect that prevents these particles from crossing the nucleus boundary or getting stuck in the space of the lamina particle mesh grid.

**Table S4. Summary of the interaction potentials between chromatin particles and nuclear landmarks and their corresponding parameter values.** See also text *Section: Chromosome-nuclear landmark interactions* for further discussion and *Section: Adam optimizer for chromosome-nuclear body interaction parameters* for details on the optimization algorithm.

| Potentials | Functional forms | Parameter values |
|---|---|---|
| Chromatin-Nucleolus | $U_{\text{C-No}}(r_{ij})$ in Eq. S26 | $\eta = 4.0$ provides a smooth transition in the tanh function for contacts. $r_c = 0.75$ reflects the minimal distances between chromatin and nucleolus beads as reflected in the excluded volume potential defined in Table S3. The interaction strength of the $i$-th chromatin bead $\alpha_i^{\text{C-No}} = P_i^{\text{N}}$, where $P_i^{\text{N}}$ is the probability for the chromatin bead $i$ to contact nucleoli as quantified by the software SPIN [24]. |
| Chromatin-Speckle | $U_{\text{C-Sp}}(r_{ij})$ in Eq. S25 | $\eta = 4.0$ and $r_c = 0.75$ were similarly determined as in $U_{\text{C-No}}(r_{ij})$. Value for the interaction strength of the $i$-th chromatin bead $\alpha_i^{\text{C-No}}$ was obtained from optimizations against SON TSA-Seq data. |
| Chromatin-Lamina | $U_{\text{C-La}}(r_{ij})$ in Eq. S24 | $\eta = 4.0$ and $r_c = 0.75$ were similarly determined as in $U_{\text{C-No}}(r_{ij})$. Value for the interaction strength of the $i$-th chromatin bead $\alpha_i^{\text{C-No}}$ was obtained from optimizations against Lamin B DamID data. The extra Lennard Jones potential was included to provide the excluded volume effect, with $\epsilon_{\text{LJ}} = 1.0$ and $r_{\text{cut}} = 0.75 \times 2^{1/6}$ as standard values. $\sigma_{\text{LJ}} = \frac{\sigma_{\text{C}} + \sigma_{\text{La}}^{\dagger}}{2} = 0.75$. |

$^{\dagger}$As mentioned in the *Section: Mapping lamina bead size to real unit*, a larger value for $\sigma_{\text{La}}$ was used here to provide a stronger excluded volume effect that prevents these particles from crossing the nucleus boundary or getting stuck in the space of the lamina particle mesh grid.

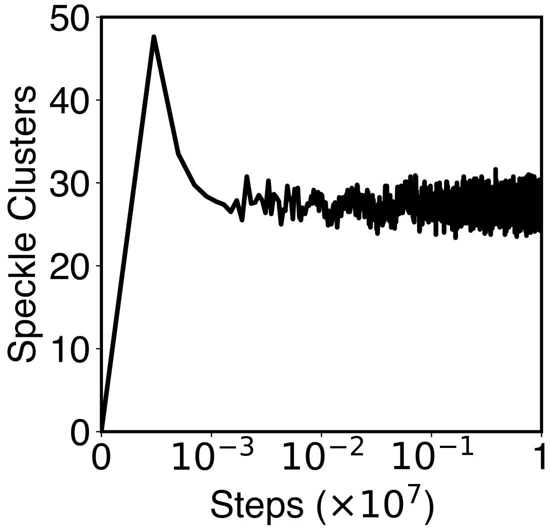

**Fig S1. The number of speckle clusters formed along a typical simulation trajectory.** This plot shows that the kinetic scheme of speckle particle exchange produces a total of $\sim 30$ speckle droplets, reproducing experimental observations. See text *Section: Speckles as phase-separated droplets undergoing chemical modifications* for more details of the kinetic scheme.

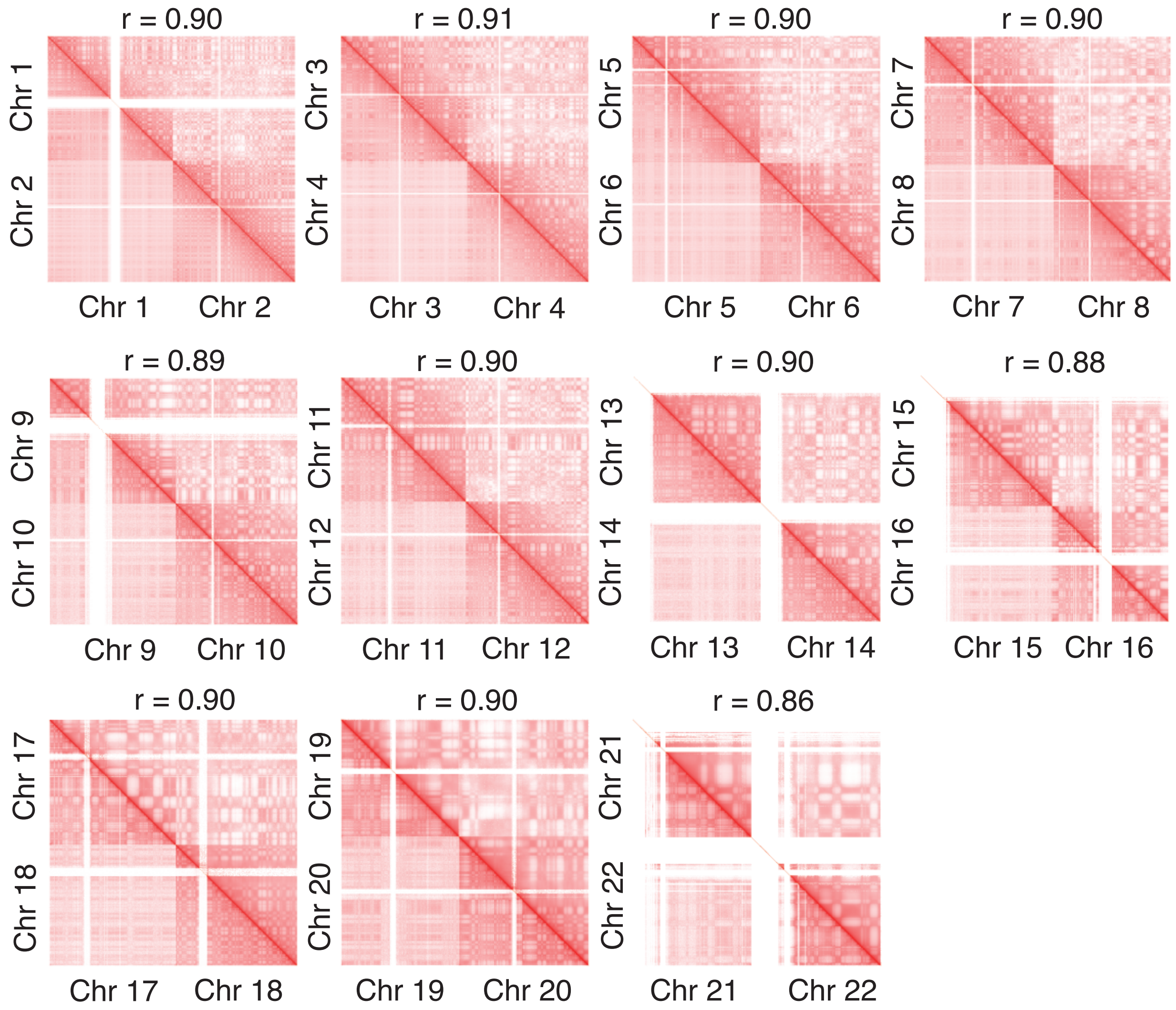

**Fig S2. Zoom-in of various regions in the contact map presented in Fig. 4 of the main text further supports the agreement between simulation and experiment.** The simulated and experimental contacts are shown in the upper and lower triangles, respectively, and the Pearson correlation coefficients $r$ between two the sets of contacts were calculated with Eq. S41.

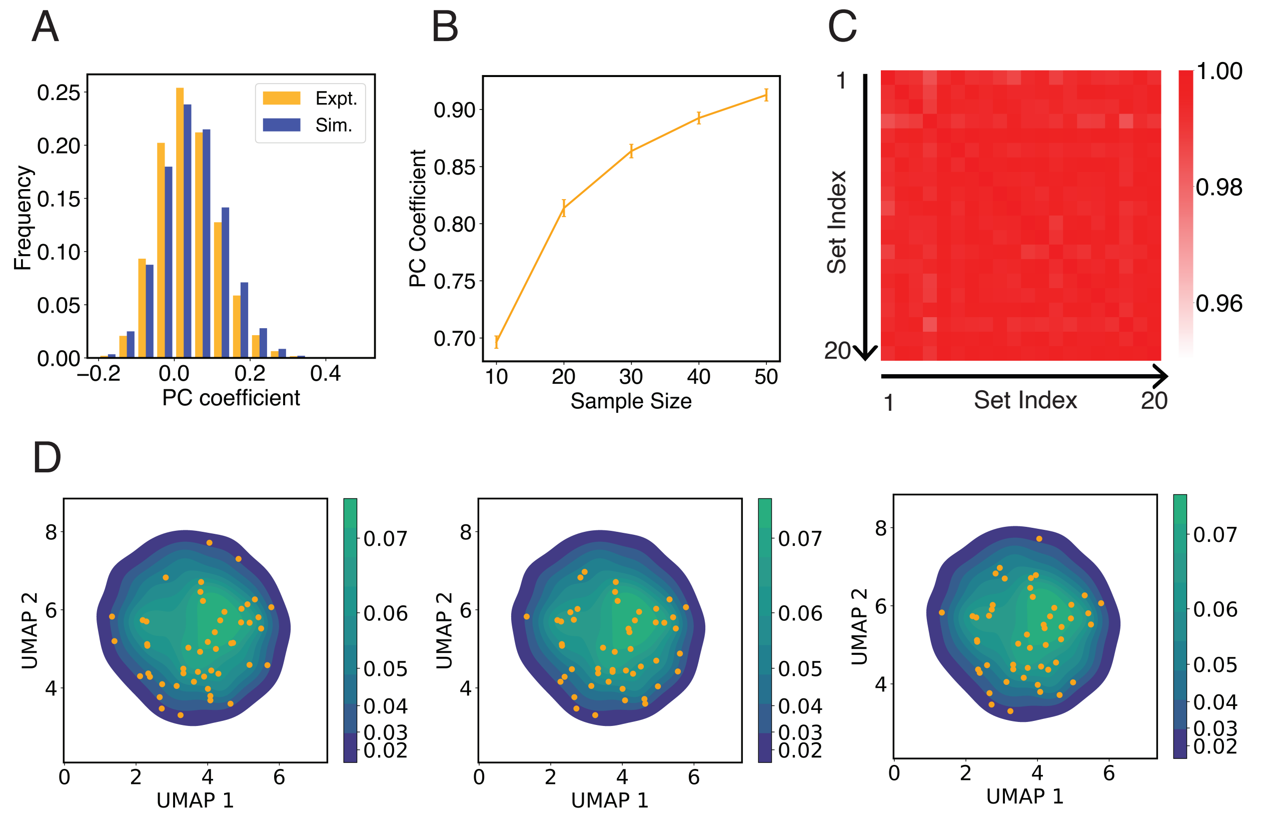

**Fig S3. Configurations used to initialize simulations capture the heterogeneity in interchromosomal contacts seen in DNA-MERFISH data.** (A) Comparison between the simulated and experimental distribution of Pearson correlation coefficients of interchromosomal contacts. Both distributions were computed using the pairwise correlations between genome structures from the respective configurational ensemble. The pairwise interchromosomal contacts include all unique haploid pairs and were computed by averaging the contact probabilities between all genomic segments from the four respective chromosomes. The simulated configuration ensemble includes 1000 unique structures prepared following the protocol outlined in *Section: Initial configurations for simulations*. The experimental ensemble includes 5455 structures reported in Ref. [31] using DNA-MERFISH. (B) Pearson correlation coefficient between average experimental and simulated pairwise interchromosomal contacts as a function of the number of independent configurations used to initialize simulations. Interchromosomal contacts are similarly defined as in part A and were computed by averaging over all configurations in the respective ensembles. To compute simulated interchromosomal contacts, we used a selection procedure as detailed in *Section: Initial configurations for simulations* to determine an ensemble of a given sample size. As the size of the ensemble increases, the resulting average interchromosomal contacts agree better with experimental results. The errorbars represent standard deviation computed from 20 independent trials. (C, D) The protocol for selecting initial configurations for simulations robustly capture the heterogeneity seen in experimental configurational ensemble. To examine the experimental structural heterogeneity, we applied Uniform Manifold Approximation and Projection (UMAP) to reduce the interchromosomal contacts into two variables following the procedure detailed in *Section: Interchromosomal contacts from DNA MERFISH data*. The plot in part C shows the Pearson correlation coefficients of 50 initial configurations represented in the UMAP variables between independent trials. The average correlation coefficient is 0.98. In part D, we show three independent initial configurations (orange dots) over the distribution of UMAP variables estimated using the experimental configurational ensemble.

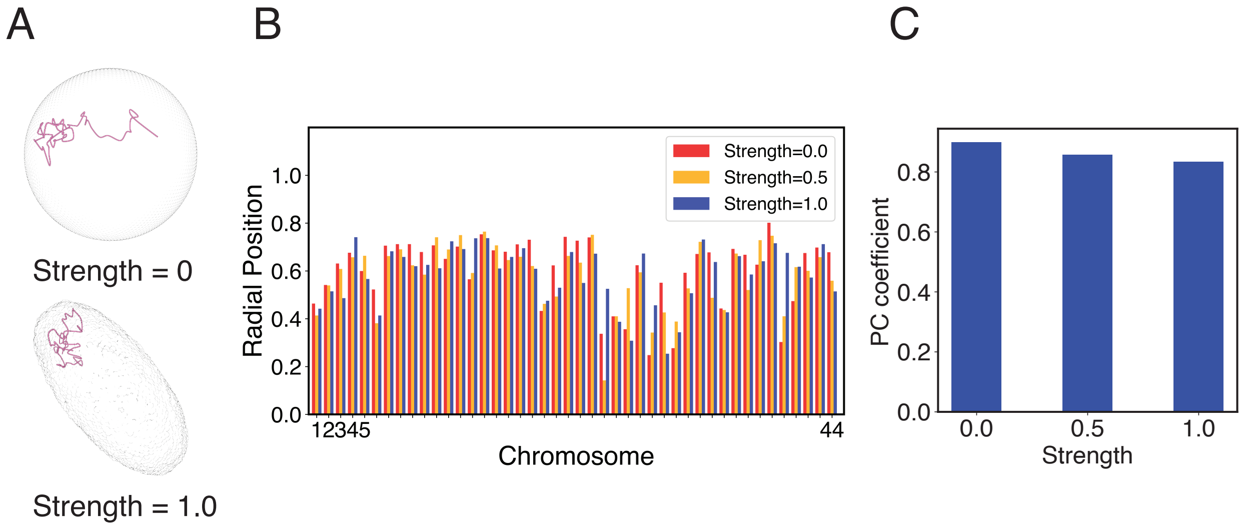

**Fig S4. Impact of nuclear deformation on normalized chromosome radial positions.** (A) The position of chromosome 21 before and after the nuclear deformation. (B) Normalized radial positions of individual diploid chromosomes at various strengths of nuclear deformation forces. (C) Pearson correlation coefficients between normalized chromosome radial positions from simulations of deformed nuclei and those from a spherical nucleus. See text *Section: Nuclear envelope deformation simulations* for simulation details.

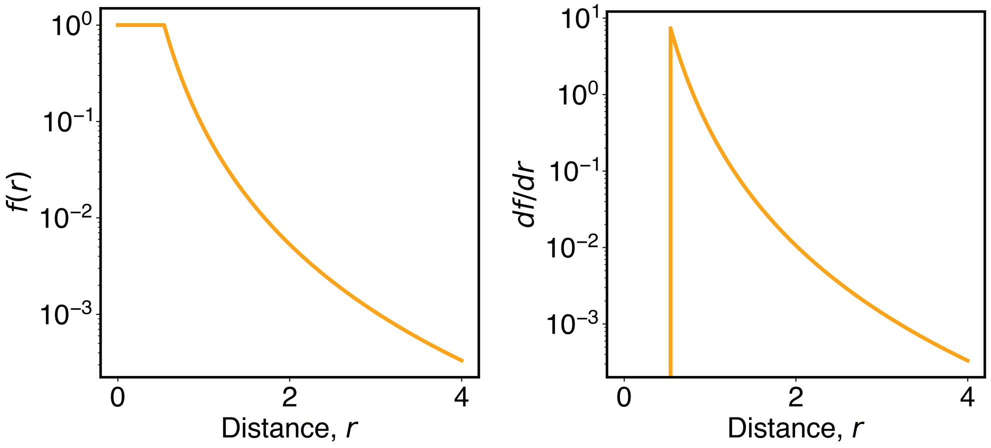

**Fig S5. The function defined in Eq. S13 smoothly switches from high to low contact probabilities.** The left and right panels plot the function and its derivative as a function of the distance, $r$.

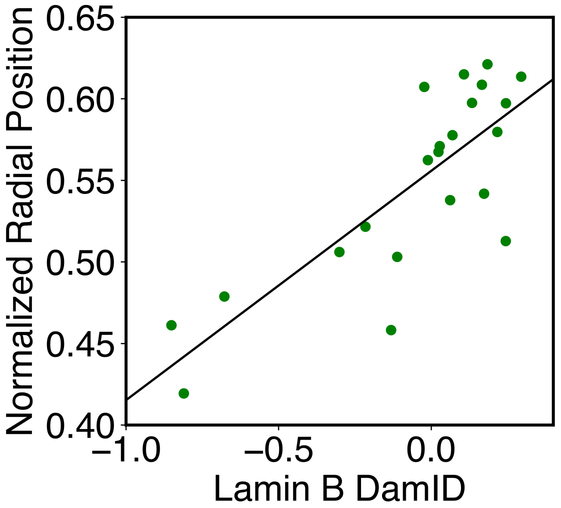

**Fig S6. Correlation between average DamID profiles of individual chromosomes with their normalized radial positions.** The normalized radial positions were determined using the average value of all cells reported from DNA MERFISH data [31]. The correlation coefficient between the two datasets is 0.8.

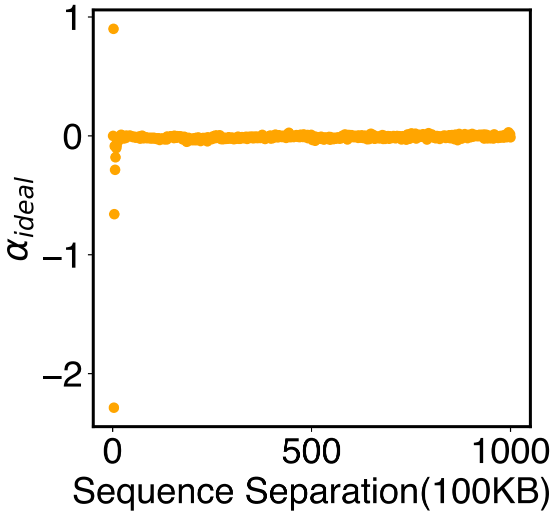

**Fig S7. Parameters of the ideal potential, $\alpha_{\text{ideal}}$, as defined in Eq. S12.** Numerical values for $\alpha_{\text{ideal}}$ are included in the software's GitHub repository.

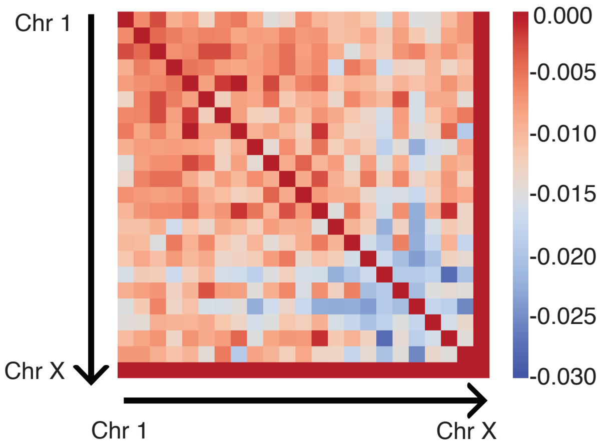

**Fig S8. Parameters of the inter potential, $\alpha_{\text{inter}}$, as defined in Eq. S15.** Numerical values for $\alpha_{\text{inter}}$ are included in the software's GitHub repository.

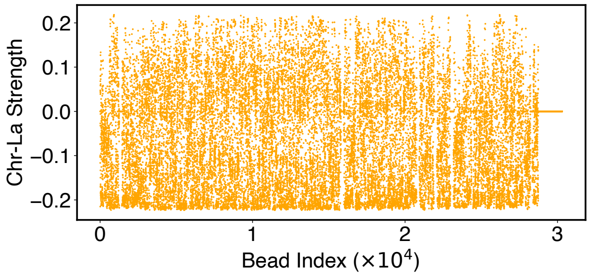

**Fig S9. Parameters of the Chromosome-Lamina potential, $\alpha^{C-La}$, as defined in Eq. S24.** Numerical values for $\alpha^{C-La}$ are included in the software's GitHub repository.

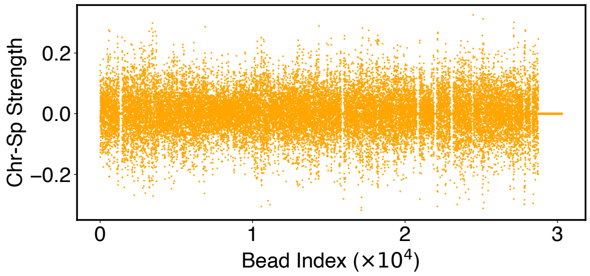

**Fig S10. Parameters of the Chromosome-Lamina potential, $\alpha^{C-Sp}$, as defined in Eq. S25.** Numerical values for $\alpha^{C-Sp}$ are included in the software's GitHub repository.

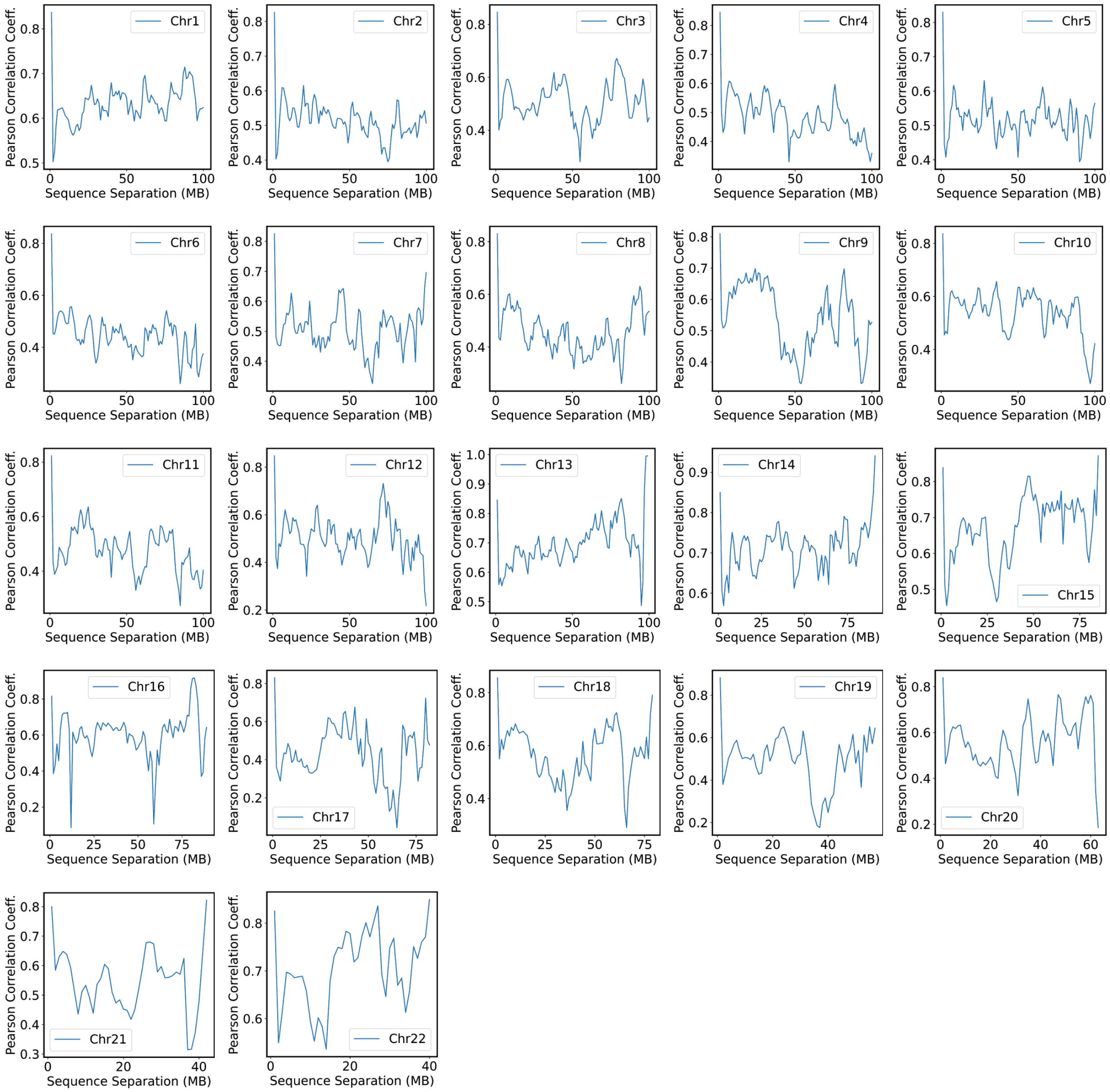

**Fig S11. Pearson correlation coefficients between experimental and simulated contact probabilities at various sequence separations within specific chromosomes.** For each chromosome, we first gathered a set of experimental contacts alongside a matching set of simulated ones for genomic pairs within a particular separation range. The Pearson correlation coefficient at the corresponding sequence separation was then determined using Eq. S41. We limited the calculations to half of the chromosome length to ensure the availability of sufficient data.

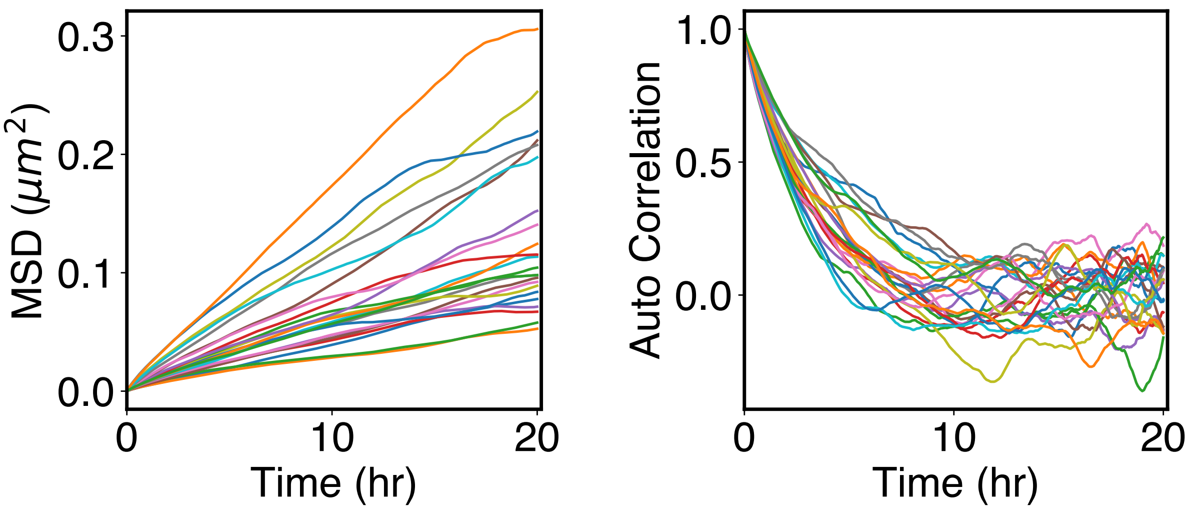

**Fig S12. Arrested kinetics of chromosome positions over the timescale of cell cycles.** (A) Mean squared displacement (MSD) of chromosome center of masses as a function of time. The MSDs are much smaller than the average size of chromosomes (see Rg values in Fig. 5A), supporting arrested dynamics. (B) Autocorrelation function of normalized chromosome radial positions as a function of time. The autocorrelation function was computed as $C(\Delta t) = \frac{\sum_{t=1}^{N}(r_t - \bar{r})(r_{t+\Delta t} - \bar{r})}{\sum_{t=1}^{N}(r_t - \bar{r})^2}$, where $t$ indexes over the trajectory frames and $\bar{r}$ is the mean position.
